# Conflict during learning reconfigures the neural representation of positive valence and approach behaviour

**DOI:** 10.1101/2023.02.03.527017

**Authors:** Laura Molina-García, Susana Colinas-Fischer, Sergio Benavides-Laconcha, Lucy Lin, Emma Clark, Neythen J. Treloar, Blanca García-Minaur-Ortíz, Chris P. Barnes, Arantza Barrios

## Abstract

Punishing and rewarding experiences can change the valence of sensory stimuli and guide animal behaviour in opposite directions, resulting in avoidance or approach. Often, however, a stimulus is encountered with both positive and negative experiences. How is such conflicting information represented in the brain and resolved into a behavioural decision? We address this question by dissecting a circuit for sexual conditioning in *C. elegans*. In this learning paradigm, an odour is conditioned with both a punishment (starvation) and a reward (mates) resulting in odour approach. We find that negative and positive experiences are both encoded by the neuropeptide PDF-1 being released from and acting on different neurons. Each experience creates a separate, parallel memory in the circuit for odour processing. This results in the sensorimotor representation of odour being different in naïve and sexually conditioned animals despite both displaying approach. Our results reveal that the positive valence of a stimulus is not represented in the activity of any single neuron class but flexibly represented within the circuit according to the experiences and predictions associated with the stimulus.

## Introduction

Animals need to constantly evaluate the stimuli they encounter to decide to approach, avoid or ignore them. The valence (i.e. the value and sign -whether good or bad-) of a stimulus is subjective and depends on genetics (and species-specific constraints), on physiological needs (or motivational state) and on previous experiences. Some stimuli, such as food odours or the pheromones of mating partners, have positive valence and are innately attractive, whereas other stimuli such as the smell of a predator are innately aversive. However, even valence that is innately assigned can be switched through learning and experience ^1, 2^. For example, the response of male fruit flies to female pheromones switches from attraction to avoidance after repeated courtship rejection ^3^, and *C. elegans* worms learn to avoid the innately attractive bacteria *Pseudomonas aeruginosa* after ingesting this pathogen ^4, 5^. How is valence appropriately and flexibly assigned? Studies in species as diverse as worms, insects, fish and mice have identified specific and segregated circuits dedicated to process positive (approach) and negative (avoidance) valence ^5–8^. Such circuits provide information about the motivational state of the animal or about the experiences associated to a stimulus to guide approach or avoidance. However, and with the exception of the gill withdrawal reflex in Aplysia ^9^, a fully mapped out circuit and path of information flow from the rewarding or punishing experience that modifies the valence assigned to a stimulus, to the behavioural decision to approach it or avoid it has not been described. Furthermore, in nature, stimuli are experienced together with positive and negative events. How is this conflict in valence assignment resolved? We address this question by dissecting the circuit mechanisms underlying sexual conditioning in *C. elegans*.

*C. elegans* worms are innately attracted to many chemical stimuli, including salt and the odour benzaldehyde which is a bacterial metabolite ^10, 11^. Innate approach however, can be switched to learned avoidance after aversive conditioning, in which worms experience the stimulus together with a punishment such as starvation ^12–14^. An additional switch in preferences has been described in males (but not hermaphrodites), for which learned avoidance can be overridden by the presence of mates during conditioning resulting in approach ^15, 16^. This form of male-specific learning, which is termed sexual conditioning, provides a paradigm to understand how conflicting rewarding and punishing experiences are integrated and resolved to assign valence and guide behaviour to a stimulus. Here, we aversively and sexually conditioned the behaviour of males to the odour benzaldehyde and asked: (1) how is information about mate-experience incorporated into the circuit for odour processing? And (2) how does this information change odour processing to guide behaviour? Specifically, because in sexual conditioning, the presence of mates overrides the behavioural consequences of aversive learning resulting in approach instead of avoidance, we postulated two alternative models. In one model, the presence of mates inhibits the formation of the aversive memory. In an alternative model, the aversive memory still forms but the presence of mates creates a parallel memory which results in approach. Through cell-specific manipulation of neuropeptide signalling, imaging of neuronal activity and behavioural analysis, we identify a circuit modulated by mate experience through the neuropeptide PDF-1 and show that two parallel memories are formed during integration of reward and punishment. The result of this is that sexually and aversively conditioned animals, although displaying opposite behaviours, have a more similar sensorimotor representation of the odour than sexually conditioned and naïve animals, which both display the same behaviour (approach). Our findings reveal that the valence of a stimulus is not represented in the activity of any single neuron class but distributed within the circuit and flexibly represented according to experience.

## Results

### Odour preferences can be sexually conditioned in males

The presence of mates has previously been shown to condition the behaviour of males to salt ^15, 16^. We asked whether sexual conditioning could modify the behaviour to stimuli of other modalities such as odours. To assess whether the responses to the odour benzaldehyde could be sexually conditioned, we first established an aversive conditioning protocol that switched the animal’s innate response to the odour from attraction to learned repulsion (aversive learning). To optimise our conditioning protocol, we exposed same-sex worm populations to the presence of odour in the absence of food for different odour concentrations and conditioning times. We determined 15% benzaldehyde for 3 hours as the optimal condition that elicits robust aversion to benzaldehyde as a result of a negative association between odour and starvation, without producing any toxicity (Supplementary Table 1). As controls, we mock conditioned animals with starvation but no odour or with odour and food, neither of which resulted in a switch in behaviour from attraction to repulsion (Fig. 1a,b). We then tested whether aversive learning could be overridden by the presence of mating partners during conditioning, as a process of sexual conditioning which has been shown to occur in males (but not hermaphrodites) during salt chemotaxis learning ^15, 16^. We observed that hermaphrodites underwent aversive learning and remained repelled by the odour regardless of whether mates (males) had been present or not during conditioning with odour and starvation (Fig. 1b). In contrast, males that were conditioned with odour, starvation and mates (hermaphrodites) did not display aversive learning and remained attracted to the odour (Fig. 1b). Together these results show that, as with salt, behavioural responses to odour can be sexually conditioned in males but not in hermaphrodites. This supports the idea that for males, who need to mate in order to reproduce, sex is a rewarding experience that can modulate the behavioural response to other environmental stimuli ^17 18^ and even override aversive learning. We next sought to identify the molecular and cellular mechanisms underlying sexual conditioning.

**Fig. 1:**
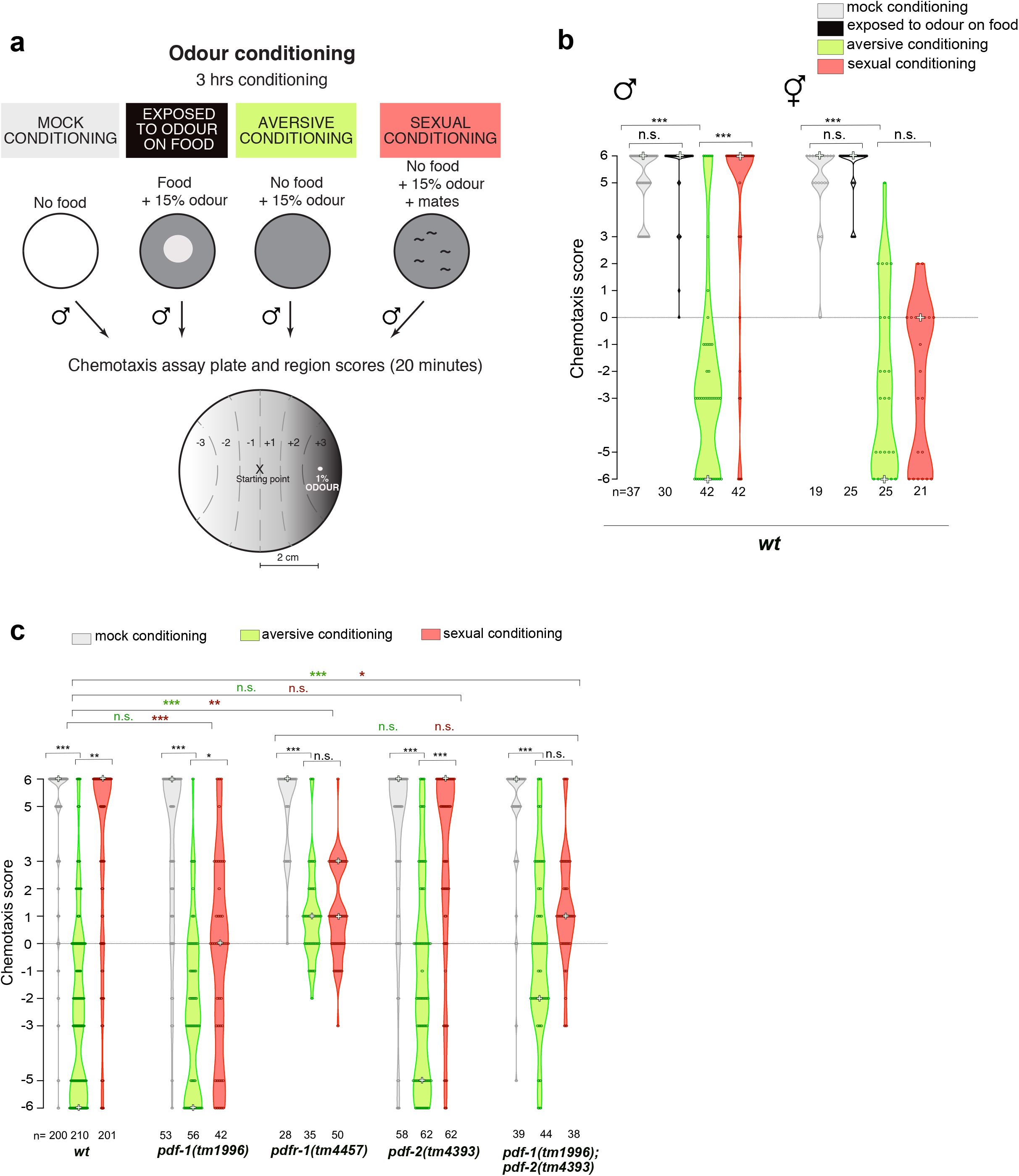
PDF neuropeptide signaling regulates aversive learning and sexual conditioning to odour. **a**. Schematic representation of the odour conditioning paradigm. Adult worms were either mock conditioned (only starved), aversively conditioned (starved in presence of 15% benzaldehyde), sexually conditioning (starved in presence of 15% benzaldehyde and mates) or exposed to 15% benzaldehyde on food for 3 hours. Individual worms were then tested for their response to a gradient of 1% benzaldehyde during 20 minutes. Their chemotaxis score was calculated based on the tracks left on the plate adding up the score of each sector through which the worm travelled. **b and c**, Chemotaxis score to benzaldehyde after conditioning. Each empty circle within a violin plot represents the chemotaxis score of an individual animal. The mode is marked with a white cross. ξ_2_ for trend analysis and Bonferroni correction was used to compare the frequency distribution of values of each sex or genotype between conditions (black asterisks) and for a condition between genotypes (green asterisks for aversive conditioning and red asterisks for sexual conditioning). *** p< 0.0001; ** p< 0.001; * p<0.01; n. s. no statistically significant difference p≥0.05. n, number of animals. In **c**, different genotypes were assayed in different days but always with a *wildtype* control (pulled together for this graph). For sexual conditioning, control and genotype of interest were conditioned in the same plate and identified through genotyping after testing. At least 3 independent experiments were done for each group. *him* alleles used were *him-8(e1489)* and *him-5 (e1490)* for b and c, respectively.

### PDF neuropeptide signalling regulates aversive learning and sexual conditioning

We have previously shown that sexual conditioning to salt is regulated by the neuropeptide Pigment Dispersing Factor 1 (PDF-1) ^16^. A second gene in the *C. elegans* genome, *pdf-2*, encodes a ligand that also acts through the PDF-1 receptor, PDFR-1 ^19, 20^. We therefore tested whether mutants for both ligands and the receptor had altered innate and learned behavioural responses to odour. To determine whether a given genotype was proficient or defective at learning, we compared the frequency distribution of chemotaxis scores between conditions. We made two comparisons: one across genotypes comparing mutant and *wildtype* worms for a given conditioning treatment and another within genotype comparing mock with aversive conditioning and aversive with sexual conditioning (see Methods). As previously reported for sexual conditioning to salt ^16^, *pdf-1(tm1996)* mutants exhibited a defect in sexual conditioning to odour compared to *wildtype* males and displayed chemotaxis scores distributed across all possible categories and mainly concentrated around 0, indicating a lack of preference for the odour instead of attraction (Fig. 1c). The defect exhibited by *pdf-1* mutants was specific for sexual conditioning as their naïve responses and aversive learning were not significantly different from control animals (Fig. 1c). To confirm the requirement of PDF signaling in sexual conditioning, we tested the responses of mutants in the receptor. Surprisingly, *pdfr-1(tm4457)* receptor mutants were defective not only in sexual conditioning but also in aversive learning. Most *pdfr-1* animals displayed a lack of preference or even some residual attraction after aversive conditioning and maintained this similar distribution after sexual conditioning (Fig. 1c). To establish whether aversive learning is mediated by another PDF ligand, we tested the responses of *pdf-2* single and *pdf-1; pdf-2* double mutants. *pdf-2 (tm4393)* mutants did not differ from *wildtype* animals in either aversive learning or sexual conditioning and exhibited clear negative and positive preference for the odour after aversive and sexual conditioning, respectively. In contrast, *pdf-1(tm1996); pdf-2(tm4393)* double mutants lacking both PDF ligands were defective in both aversive and sexual conditioning learning, like *pdfr-1* receptor mutants (Fig. 1c). Altogether these data show that both PDF-1 and PDF-2 are sufficient for aversive learning but only PDF-1 is required to modulate odour preferences after sexual conditioning.

### PDF signalling acts on different neurons to drive odour avoidance or approach after learning

Next we asked which cells respond to PDF-1 neuromodulation to effect sexual conditioning. We generated rescue constructs that express *wildtype pdfr-1* cDNA in *pdfr-1* mutant animals using two previously reported *pdfr-1* promoter sequences which drive expression in different groups of neurons ^21, 22^(Fig 2a,b). The proximal promoter was used to drive expression in neurons and body muscle ^21^ (Fig. 2b). The distal promoter was used to drive conditional expression of the receptor upon recombination with different Cre lines ^22^ (Fig. 2b).

**Fig. 2:**
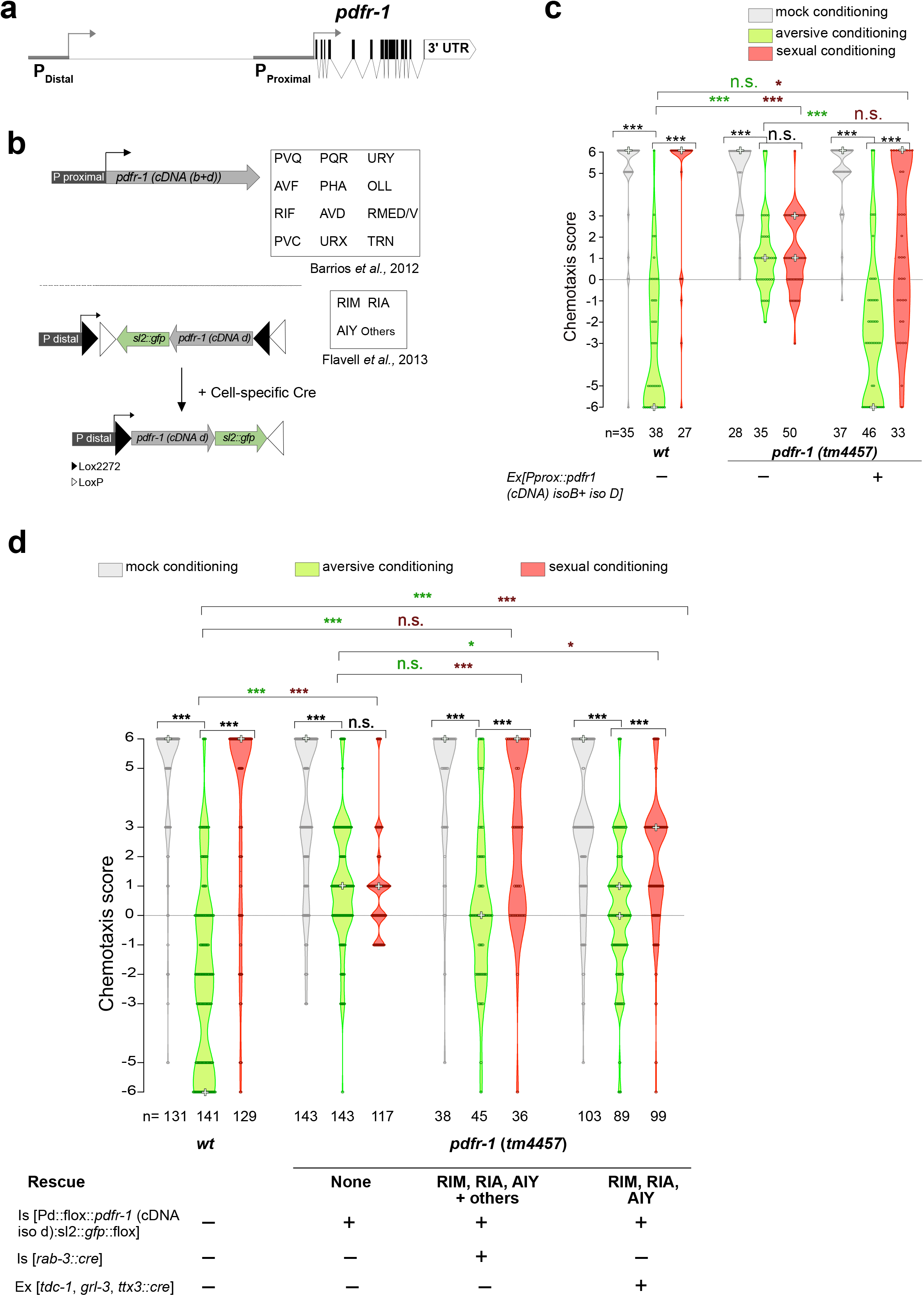
PDFR-1 is required in different neurons for learned approach and learned avoidance. **a**, Genomic locus of the *pdfr-1* gene. Promoter regions are depicted as grey bars and exons as black boxes. Arrows indicate transcription from either promoter. **b**, Schematic representation of the DNA constructs used to rescue expression of the *pdfr-1* gene. Expression of the *pdfr-1* sense mRNA from the distal promoter (bottom) relies on the intersectional expression of Cre recombinase in the cells of interest. The list of neurons in which the distal and proximal promoters drive expression is shown. **c and d**, Chemotaxis score to benzaldehyde after conditioning. Each empty circle within a violin plot represents the chemotaxis score of an individual animal. The mode is marked with a white cross. ξ_2_ for trend analysis and Bonferroni correction was used to compare the frequency distribution of values of each genotype between conditions (black asterisks) and for a condition between genotypes (green asterisks for aversive conditioning and red asterisks for sexual conditioning). *** p< 0.0001; ** p< 0.001; * p<0.01; n. s. no statistically significant difference p≥0.05. n, number of animals. *pdfr-1(tm4457)* data in **2c** is the same as in **1c**. For sexual conditioning, controls (*wildtype* and mutant) and rescued animals were conditioned in the same plate and identified by genotyping or through visualisation of the fluorescent transgene after testing. At least 3 independent experiments were done for each group. *wt* background is *him-5(e1490)*.

Expression of *pdfr-1* under the proximal promoter fully rescued odour avoidance in *pdfr-1* mutants after aversive learning (Fig. 2c). After sexual conditioning, transgene-carrying mutant worms displayed a frequency distribution of chemotaxis scores that was not significantly different from that of *pdfr-1* mutants. However, their scores were shifted towards more positive values compared to those displayed after aversive learning, indicating a loss of aversion and therefore, a partial rescue of sexual conditioning (Fig. 2c). Restoring expression of *pdfr-1* from the distal promoter with a panneuronal Cre line fully rescued odour attraction in *pdfr-1* mutants after sexual conditioning, with a distribution of chemotaxis scores no significantly different from that of *wildtype* worms (Fig. 2d). This construct however, did not rescue odour avoidance after aversive conditioning and worms displayed a lack of preference, similar to *pdfr-1* mutants (Fig. 2d). Together, these data suggest that the PDFR-1 receptor is required in different neurons to drive either aversion or attraction to the odour after learning.

The distal but not the proximal *pdfr-1* promoter has been shown to drive expression in the interneurons RIM, RIA and AIY ^22^. These neurons are part of the circuit for odour processing ^23, 24^ and have been implicated in naïve and learnt behavioural responses to other sensory stimuli ^5, 25, 26^. We therefore tested whether expression of *pdfr-1* only in these 3 classes of neurons could rescue odour attraction after sexual conditioning in *pdfr-1* mutants. For this, we used the *tdc-1* ^22^, *glr-3*^22, 27^ and *ttx-3* ^28^ promoters to drive expression of Cre recombinase in RIM, RIA and AIY, respectively. Expression of *pdfr-1* in RIM, RIA and AIY together partially rescued odour attraction after sexual conditioning, shifting the chemotaxis scores of *pdfr-1 (tm4457)* males towards more positive values (Fig. 2d). After aversive learning, although the distribution of chemotaxis scores of transgene-carrying *pdfr-1* mutants was significantly different from that of mutants without the transgene, scores remained distributed across all possible categories and mainly concentrated around 0, indicating a lack of preference for the odour (Fig. 2d). Altogether, these results indicate that odour attraction after sexual conditioning but not odour avoidance after aversive learning, requires modulation of RIM, RIA and AIY by PDF-1.

### PDF-1 is required in the interneurons AVB and MCM to mediate sexual conditioning

We next set out to identify the neurons that are important sources of PDF-1 to mediate odour attraction after sexual conditioning. During sexual conditioning, the information about mate experience must be incorporated into the circuit for odour processing ^15^. We reasoned that neurons that receive inputs from circuits dedicated to sensing mate cues may be good candidates to test. We identified two classes of *pdf-1*-expressing neurons with such connectivity: the AVB interneurons, which are present in both sexes, and the male-specific MCM interneurons, which we have previously shown to be required for sexual conditioning to salt ^16, 17, 29–32^ (Fig. 3a,b).

**Fig. 3:**
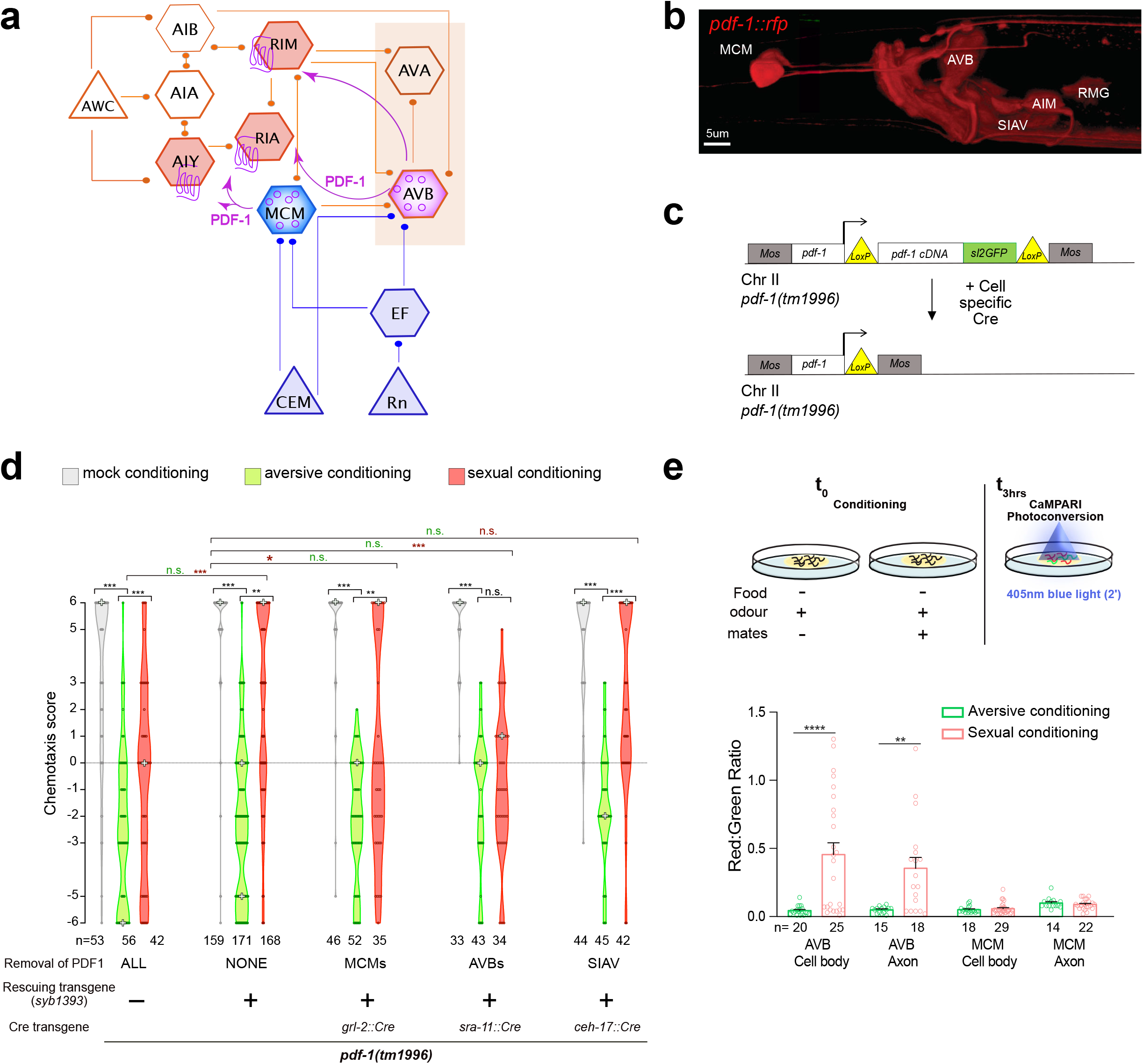
The MCM and AVB interneurons are a source of PDF-1 for sexual conditioning. **a**, Diagram of the circuit for odour processing in males. Triangles, hexagons and circles represent sensory neurons, interneurons and PDF-1 neuropeptide, respectively. Synaptic connections are depicted as lines and peptidergic communication as arrows. Male specific neurons and PDF-1 target neurons are shown in blue and pink, respectively. The names of the different neurons within the circuit are indicated. **b**, Lateral view (anterior to the left) of the anterior-most part of a male expressing a *pdf-1::rfp* reporter transgene. Some neurons are labeled with their names. **c**, scheme of the excisable *pdf-1* transgene integrated as single copy in a MosCI site in chromosome II. **d**, Chemotaxis score to benzaldehyde after conditioning. Different genotypes were assayed in different days but always in parallel to a positive control (*him-5 (e1490) V; pdf-1(tm1996)III;syb1393*). Data for *pdf-1(tm1996)* mutants is the same as in Fig 1c. For sexual conditioning, controls and genotype of interest were conditioned in the same plate and identified using fluorescence microscopy after testing. At least 3 independent experiments were done for each genotype. All strains were in *him-5(e1490)* background. For **a** and **c**, each empty circle within a violin plot represents the chemotaxis score of an individual animal. The mode is marked with a white cross. ξ_2_ for trend analysis and Bonferroni correction was used to compare the frequency distribution of values of each genotype between conditions (black asterisks) and for a condition between genotypes and temperatures (green asterisks for aversive conditioning and red asterisks for sexual conditioning). *** p< 0.0001; ** p< 0.001; * p<0.01; n. s. no statistically significant difference p≥0.05. n, number of animals. **e**, Top, schematic of the CaMPARI photoconversion experiment. **e**, Bottom, measurements of CaMPARI photoconversion in AVB and MCM interneurons after conditioning. Each circle corresponds to a neuron. Two tail Mann-Whitney U test was used to compare conditions. Black asterisks show statistically significant difference between different conditioning protocols ****p<0.0001; ** p<0.01. Error bars represent SEM. n, number of neurons.

To determine whether *pdf-1* is required in the MCM and AVB interneurons for sexual conditioning, we generated a strain that contained a single copy insertion of a rescue construct for *pdf-1* flanked by Lox P sites to enable cell-type specific loss-of-function of *pdf-1* ^22^ (Fig. 3c). This transgene fully rescued the sexual conditioning defects of *pdf-1(tm1996)* mutants (Fig. 3d). We then excised *pdf-1* from specific subsets of neurons using different Cre recombinase strains (Supplementary Table 2). Removing *pdf-1* from the MCM interneurons resulted in a significant shift in the distribution of chemotaxis scores towards more negative values and across all categories (Fig. 3d). Removing *pdf-1* from AVB interneurons resulted in males that failed to undergo sexual conditioning and displayed a similar distribution of chemotaxis scores after sexual conditioning and after aversive learning (Fig. 3d). In contrast, removing *pdf-1* from the SIAV interneurons, which do not receive any synaptic input from mate-sensing circuits ^29^, did not result in any defects in naïve or learned odour preferences (Fig. 3d). These results demonstrate that not all *pdf-1-*expressing neurons are relevant sources of neuropeptide release during learning and that sexual conditioning requires *pdf-1* from both male-specific MCM and sex-shared AVB interneurons.

Removing *pdf-1* from MCMs or AVBs resulted in defects specific to sexual conditioning while the naïve and aversively conditioned responses to odour were unperturbed. However, PDF-1 also regulates aversive learning, a role that is masked by the presence of PDF-2 (Fig. 1c). To test whether MCM and AVB neurons are also relevant sources of PDF-1 for aversive learning, we carried out the same intersectional strategy of *pdf-1* removal in a *pdf-1(tm1996); pdf-2(tm4393)* double mutant background. We found that removing *pdf-1* from either MCMs or AVBs had no effect on the odour avoidance responses of aversively conditioned animals (Extended Data Fig. 1). Together, these data show that PDF-1 conveys different information to the circuit for odour processing depending on its source of release.

### AVB interneurons are activated by mate sensation

Having established that the MCM and AVB interneurons regulate sexual conditioning through the release of *pdf-1* and that they receive connectivity from circuits dedicated to sensing mates, we hypothesised that these interneurons may be activated by mate sensation during conditioning. To test this, we measured changes in intracellular Ca^2+^, as a proxy for neuronal activation, during conditioning. We used the *pdf-1* promoter to drive expression of the Calcium Modulated PhotoActivatable Ratiometric Indicator CaMPARI. This Ca^2+^ indicator irreversibly photoconverts from green to red upon blue light stimulation at high levels of intracellular Ca^2+ 33^. We measured and compared levels of photoconversion (i.e. red:green ratio) in the somas and processes of AVB and MCM neurons in animals that had been conditioned with odour and lack of food in the absence (aversively conditioned) or presence (sexually conditioned) of mates (Fig. 3e). Consistent with the hypothesis that mate sensation activates the AVB neurons, we saw high levels of photoconversion in both somas and processes of AVB neurons in sexually conditioned but not aversively conditioned males (Fig. 3e). In contrast, and opposite to our expectations, we did not observe an increase in Ca^2+^ levels in MCM neurons, which displayed low and similar levels of photoconversion in animals that had been conditioned in the absence or presence of mates (Fig. 3e).

Altogether, the experiments presented here demonstrate that AVB interneurons are stimulated by mate sensation and become active during conditioning. PDF-1 neuropeptide released from AVB neurons during conditioning modulates the circuit for odour processing to switch subsequent behavioural responses to odour from repulsion to attraction. Furthermore, our results suggest that although the MCM interneurons are required for the modulation of odour preferences by mate sensation, their activation may not be reflected as changes in intracellular Ca^2+^ levels.

### The aversive memory forms in both aversively and sexually conditioned animals

Our data shows that, during conditioning, starvation and mate sensation can act as punishing and rewarding experiences, respectively, and that they modulate future behavioural responses to the same odour stimulus in opposite directions, resulting in avoidance or approach. However, during sexual conditioning, when the odour is presented with both starvation and mates, the behavioural response is not something intermediate like lack of preference but it is approach. This raises the question of how does mate experience override the behavioural consequences of aversive learning? We envisaged two possible alternative models (Fig. 4a). In model 1, the presence of mates inhibits the formation of the aversive memory in which odour predicts starvation. In this model, only the appetitive (positive) memory in which odour predicts mates, is formed. Alternatively, in model 2, the presence of mates creates a parallel memory that co-exists with the aversive memory, which is also formed. Because memories can be identified as biophysical changes in the brain, to distinguish between the two models, we monitored and compared neuronal activity upon odour stimulation and other cellular changes within the circuit for odour processing after mock, aversive, and sexual conditioning. The expectation was that in aversively conditioned animals we would observe evidence of an aversive memory having formed as changes in neural activity within the circuit compared to naïve (mock conditioned) animals. If model 1 is correct, the changes observed in aversively conditioned animals would be absent in sexually conditioned animals. In contrast, if model 2 is correct, the changes observed after aversive learning would remain after sexual conditioning.

**Fig. 4:**
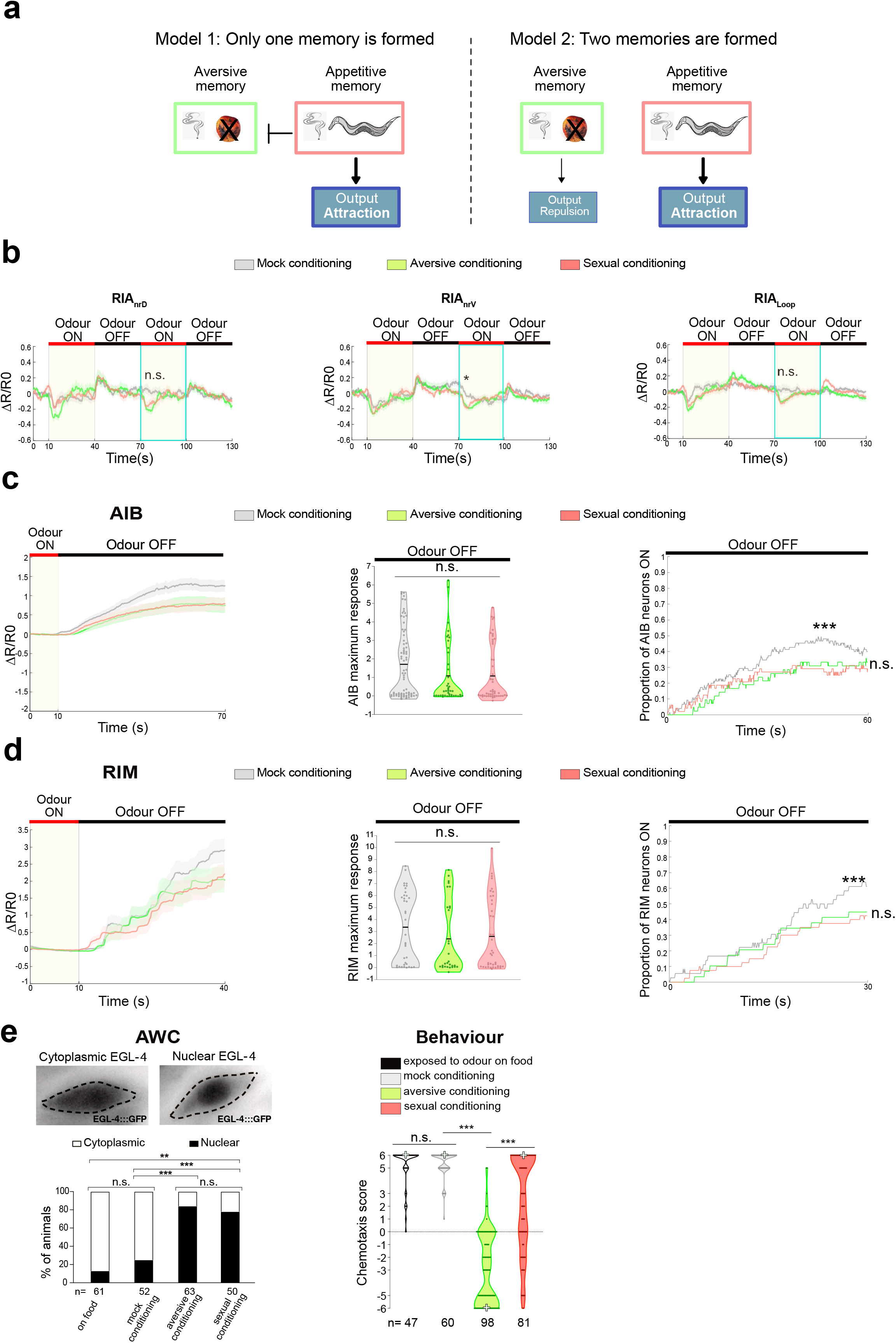
The aversive memory is also formed during sexual conditioning. **a**, Schematic representation of two proposed models for how punishment and reward are integrated during sexual conditioning. **b**, Average baseline-adjusted traces of GCaMP6f/RFP fluorescence ratios in dorsal nerve ring (nr_d_), ventral nerve ring (nr_v_) and loop domains of RIA interneurons upon odour addition and removal after different conditioning protocols. Turquoise rectangles frame the odour stimulation step in which mock-conditioned animals show adaptation. Non-parametric ANOVA was used to compare the responses of the 3 RIA domains between conditions during the second odour exposure (70-75 seconds). * p<0,01; n.s. no statistically significant difference p≥0.05. 32 neurons were analysed for mock, 23 for aversive, and 24 for sexual conditioning for RIA nrd. 26 neurons were analysed for mock, 24 for aversive, and 31 for sexual conditioning for RIA nrv. 28 neurons were analysed for mock, 23 for aversive, and 26 for sexual conditioning for RIA loop. **c**, AIB activity. Left, average baseline-adjusted traces of GCaMP6f/RFP fluorescence ratios in AIB neurons. Only neurons with low activity (normalised fluorescence R-Rmin/Rmax <0.5) before odour removal were selected. 75 neurons were analysed for mock, 42 for aversive, and 48 for sexual conditioning. Middle, maximum activity in AIB neurons during the period of odour removal (60 sec). Each dot represents one neuron. Average is indicated with a black line. Non-parametric ANOVA test was used to compare conditions. n. s. no statistically significant difference p≥0.05. Right panel, proportion of AIB neurons that became activated at each time point during odour removal. Non-parametric ANOVA test was used to compare conditions. *** p< 0.0001. **d**, RIM activity. Left, average baseline-adjusted traces of GCaMP6f/RFP fluorescence ratios in the axon of RIM neurons. Only neurons with low activity (normalised fluorescence R-Rmin/Rmax <0.5) before odour removal were selected. 36 neurons were analysed for mock, 29 for aversive, and 40 for sexual conditioning. Middle, maximum activity in the axon of RIM neurons during the period of odour removal (60 sec). Each dot represents one neuron. Black lines represent the average. Non-parametric ANOVA test was used to compare conditions. n. s. no statistically significant difference p≥0.05. Right panel, proportion of RIM neurons that became activated at each time point during odour removal. Non-parametric ANOVA test was used to compare conditions. *** p< 0.0001, n. s. no statistically significant difference p≥0.05. For b-e, at least three independent experiments were done for each condition. **e**, Top left, images of EGL4::GFP localisation in the cytoplasm and the nucleus of AWC neurons. Dashed lines indicate the edge of the AWC sensory neuron. Bottom left, percentage of animals with nuclear and cytoplasmic EGL-4 in AWC after being exposed to the different conditions. ξ_2_test with Bonferroni correction for multiple comparisons was used to compare conditions. Right panel, chemotaxis score of animals carrying the *odr-3::egl-4::gfp* transgene. Each empty circle within violin plots represents the chemotaxis score of an individual animal. The mode is indicated with a white cross. ξ_2_ for trend analysis and Bonferroni correction was used to compare the frequency distribution of values of each group between conditions. Black asterisks show statistically significant difference between different types of conditioning. *** p< 0.0001; n. s. no statistically significant difference p≥0.05.

We measured neuronal activity with the genetically-encoded Ca^2+^ indicator GCaMP6f upon odour presentation in an olfactory chip ^34, 35^. We monitored Ca^2+^ levels in 4 classes of interneurons (AIY, AIB, RIM and RIA) which act postsynaptically to the odour-sensing neuron AWC (Fig 3a). These interneurons control navigation in an odour gradient by driving forward movement (AIY) or reorientations (AIB, RIM, RIA) ^23, 24, 36–38^; Moreover, our data shows that at least three of them (AIY, RIM and RIA) receive PDF-1 neuromodulation during sexual conditioning (Fig. 2d). RIA interneurons have three axonal domains (dorsal (nr_d_), ventral (nr_v_) and sensory loop) that respond in a coordinated manner to odour presentation and removal ^37^. During a first round of stimulation, we observed a decrease in Ca^2+^ upon odour presentation and an increase upon odour removal in naïve and conditioned animals (Fig. 4b). However, RIA responses in naïve animals became rapidly adapted and we did not observe a fall in Ca^2+^ levels upon second odour stimulation (Fig. 4b). In contrast, we did not observe adaptation in aversively or sexually conditioned animals (Fig. 4b) revealing a memory trace linked to conditioning.

In the absence of odour, AIB and RIM neurons have been shown to alternate between high and low activity states ^24^ (Extended Data Fig. 2a, b). Addition of odour inhibits AIB and RIM, and odour removal activates them ^23, 24^. The responses of AIB and RIM to odour addition and removal are probabilistic and are constrained by whether the neuron is already in a high or low state of activity ^24^. To examine the responses of AIB and RIM in naïve and conditioned animals, we classified the neurons according to their high or low activity state at the point of odour removal (see Methods). Similar to what has previously been shown for well-fed worms, we observed activation upon odour removal in naïve/mock-conditioned males (Fig. 4c). In aversively conditioned animals, in contrast, AIB activation upon odour removal was reduced (Fig. 4c). We observed a trend (albeit not significant) towards lower amplitude responses as well as a significant reduction in the number of neurons that responded to odour removal (Fig. 4c). Importantly, a similar reduction in AIB activity and number of neurons that became activated upon odour removal was observed in sexually conditioned animals (Fig. 4c) consistent with the idea that the aversive memory is also formed during sexual conditioning (model 2). Similar to what we observed for AIB cell bodies, the axon of RIM interneurons became depolarised upon odour removal in naïve/mock-conditioned males and, although we did not observe a change in the magnitude of the responses, the number of RIM neurons that responded to odour removal was significantly reduced in both aversively and sexually conditioned animals (Fig. 4d). To rule out the possibility that in conditioned animals there was a general lower probability of AIB and RIM being in a high activity state, we quantified the number of neurons in a high activity state during the baseline period, prior to odour exposure.

The proportion of neurons in a high activity state was similar in conditioned and naïve/mock animals (Extended Data Figure 2c, d), therefore demonstrating that the reduction we observed in conditioned animals was specific to odor removal. Together, these results show that the memory formed during aversive learning, by pairing odour with starvation and characterised by a significant reduction of responses to odour removal in AIB and RIM, is also formed during sexual conditioning.

We sought to strengthen our observations that aversive memories are still formed in sexually conditioned animals. A cellular signature of aversive odour learning is the nuclear translocation of the cGMP-dependent protein kinase PKG/EGL-4 in the odour-sensing neuron AWC ^39, 40^. To measure the cellular localisation of PKG/EGL-4 we used a transgene driving expression of a GFP-tagged EGL-4 in AWC ^39^. As previously reported for butanone aversive learning ^39, 40^, we observed a significant increase (from 25% to 84.1 %) in the number of AWC neurons with EGL-4 nuclear localisation in animals aversively conditioned to benzaldehyde compared to naïve/mock-conditioned animals (Fig. 4e). Consistent with the idea that the aversive memory is formed also during sexual conditioning, we observed a similar proportion (76.5%) of AWC neurons with EGL-4 nuclear localization in sexually conditioned animals, despite these animals displaying attraction to the odour (Fig. 4e). Importantly, EGL-4 nuclear localisation was a consequence of pairing odour with starvation rather than a consequence of odour exposure during conditioning because conditioning animals with odour while on food did not result in EGL-4 nuclear translocation (Fig. 4e). Altogether, these data are against model 1, which proposes that the presence of mates during conditioning inhibits the formation of the aversive memory, and consistent with model 2.

### Reward and punishment form two parallel memories during sexual conditioning

Having established that the aversive memory still forms in sexually conditioned animals, we next asked whether a rewarding memory associated with mate experience also forms in the circuit for odour processing during sexual conditioning. We imaged neural activity in AIY neurons, which drive opposite behavior to RIA, AIB and RIM, inhibiting reorientations and promoting forward movement ^38, 41, 42^. The ventral domain (zone 2) of the axon of AIY interneurons has been shown to respond to odour stimulation with a rise in Ca^2+^ levels ^23^. Consistent with this, we observed a raise in Ca^2+^ in AIY axons of naïve/mock-conditioned animals upon odour presentation (Fig. 5a, b). In contrast to what we had observed for RIA, AIB and RIM, the responses of AIY did not change after aversive learning and AIY axons displayed a raise in Ca^2+^ levels similar to those of naïve/mock-conditioned animals (Fig. 5a, b). In sexually conditioned animals however, we observed a qualitative change in the responses of AIY. A significant proportion of animals (39%) displayed a delayed (maximum peak at least 15 seconds after odor exposure) increase in Ca^2+^ of high magnitude (R-Rmin/Rmax > 0.4) following the initial Ca^2+^ increase of lower magnitude upon odour presentation (Fig. 5a, b). This type of response, which we termed type 2 response (Extended Data Fig. 3a) was significantly less represented in naïve/mock-conditioned (21%) and aversively conditioned (17%) animals (Fig. 5b, c), suggesting that it may be a signature of sexual reward. As shown above, sexual reward is conveyed by the neuropeptide PDF-1 and AIY responds to PDF-1 to drive odour approach after sexual conditioning (Fig. 2d). We therefore looked at the effect that loss-of-function of *pdf-1* had on the Ca^2+^ response of AIY in sexually conditioned animals. In sexually conditioned *pdf-1 (tm1996)* mutant males, the proportion of AIY type 2 responses was significantly reduced to levels similar to those of mock and aversively conditioned animals (Fig. 5a-c and Extended Data Fig 3). These results demonstrate that sexual conditioning produces specific changes in the responses of AIY to odour presentation which are dependent on PDF-1 neuromodulation, suggesting that these changes may be important for odour approach after sexual conditioning.

**Fig. 5:**
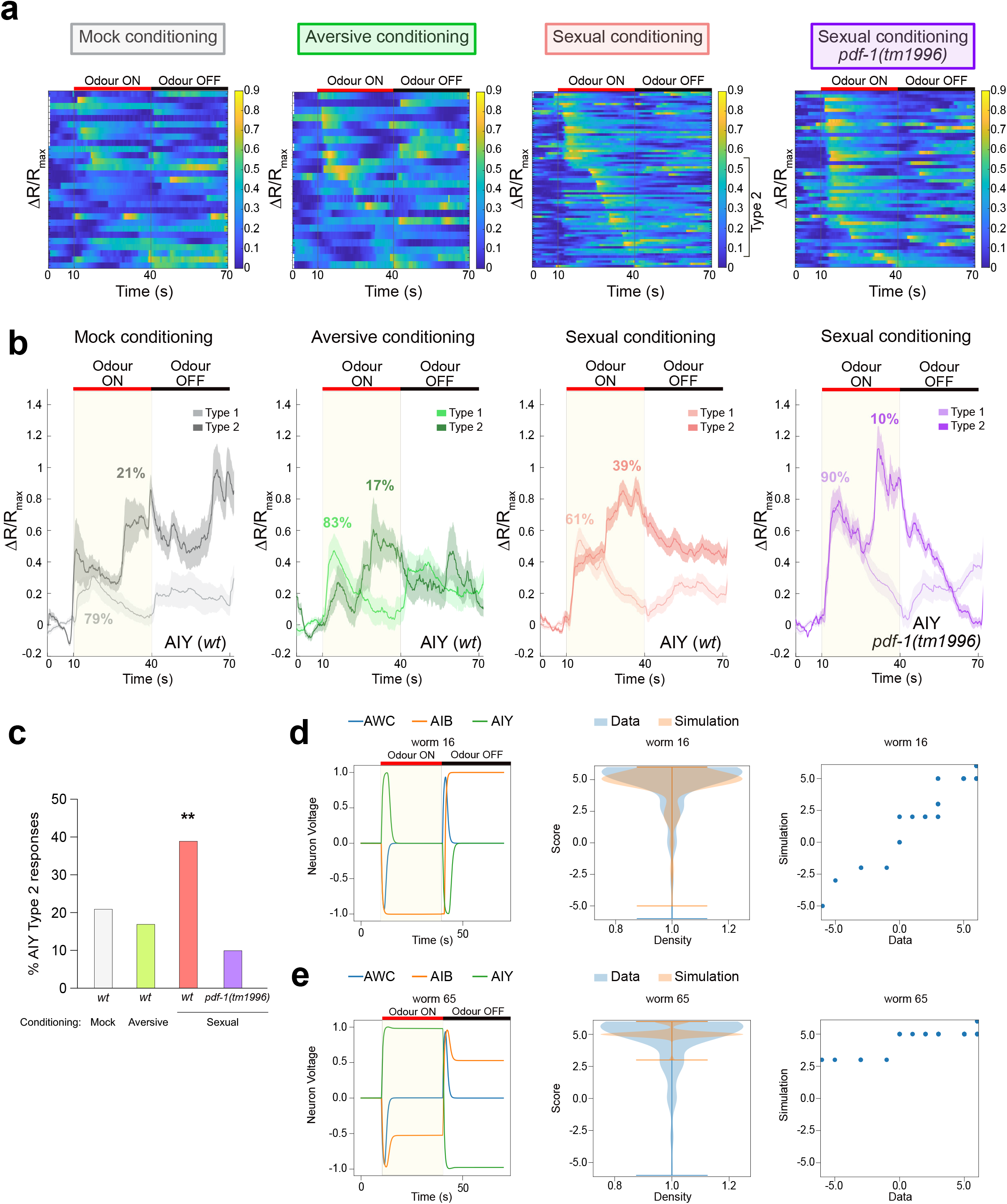
Sexual conditioning changes the neural representation of odour in AIY neurons in a PDF-1-dependent manner. **a**, Heatmaps of normalized GCaMP6f/RFP fluorescence ratios (R-R_min_)/R_max_) in the axon of AIY interneurons upon odour exposure after conditioning in *wild type* and *pdf-1* mutants. Each row represents an individual neuron. Neurons are sorted according to time to maximum activation. Type 2 responses are indicated in the sexual conditioning group, where they are most evident and frequent. **b**, average normalised traces of GCaMP6f/RFP fluorescence ratios (R-R_min_)/R_max_) in the axons of AIY neurons. Type 1 and type 2 responses have been plotted separately for each condition and their frequency indicated. 29 neurons were analysed for mock, 24 for aversive, and 75 for sexual conditioning for AIY *wild type*. 35 neurons were analysed for mock, 36 for aversive, and 50 for sexual conditioning for AIY *pdf-1(tm1996)*. **c**, Proportion of type 2 responses in the axons of AIY neurons in *wild type* and *pdf-1 (tm1996)* mutants after conditioning. ξ_2_ test was used to compare groups. ** p< 0.001. At least 3 independent experiments were performed for each group. **d**, simulated neural activity (left panel) and worm behaviour (middle and right panels) of a population with transient activity in AIY and sustained activity in AIB. Right panel is a QQ plot comparing the probability distribution of chemotaxis scores in the simulated and observed worm population. **e**, same as d but for a population with sustained activity in AIY and reduced activity in AIB. For **d** and **e**, bars within violin plots represent the 10 and 90 percentile of each distribution.

### Sustained activity in AIY can lead to robust attraction despite reduced activity in AIB

Our data demonstrates that during sexual conditioning, two distinct memories, one related to aversive learning and one related to sexual reward, form in parallel and compete for behavioural expression. Combined with knowledge from previous studies of what the motor output of the neurons in the odour circuit is, our findings allow us to build a partial picture of how the memories created after sexual conditioning may drive approach. The reduction of AIB and RIM activity upon odour downsteps will result in a reduction in the probability of reorientation when the animal is moving away from the odour source. While lack of reorientation may not account for the observed odour avoidance of aversively conditioned animals, it will result in reduced attraction and lack of preference ^38, 40^. The inability to reorient when moving away from the odour may be compensated in sexually conditioned animals by the sustained and high activity of AIY upon odour upsteps, which will result in longer forward movement when the animal is moving towards the odour source. To test the hypothesis that sustained type 2 responses in AIY can lead to robust attraction, we built a simplified ordinary differential equation model of the odourtaxis circuit including only the sensory neuron (AWC) and the first layer of postsynaptic interneurons (AIB, AIA and AIY). We used behavioural data from mock conditioned animals to fit the synaptic weights of a population of 100 simulated worms using an evolutionary algorithm over 100 generations (Extended Data Fig. 4). We then examined the simulated activity of AIY and AIB neurons in the final generation, which clustered into different patterns of neural activity that correlated with different populations of chemotaxis scores (Extended Data Fig. 5-7). Although none of the simulated worms fully recapitulated the observed dynamics of activity in AIY and AIB, a good approximation of naïve/mock responses showing transient AIY activity upon odour presentation and prolonged AIB activity after odour removal was present in 20% of the population. The simulated worm behavior most frequently linked to this neural activity pattern was attraction, with scattered scores centered around +3 (Fig. 5d; Extended Data Fig. 5-7). The model also produced simulated activity of AIY that approximated type 2 responses, sustained instead of transient, like the ones observed in sexually conditioned animals, in 45% of the population. Some of these were in the context of naïve/mock-like AIB responses and some (9%) were in the context of reduced AIB activity. In the latter group, the combination of sustained, type 2-like responses in AIY and reduced, aversive memory-like activity in AIB correlated with chemotaxis scores shifted towards higher positive values all above +3 and/or concentrated at +5 in 55% of the cases (Fig. 5e; Extended Data Fig. 5-7). Altogether, the simulations produced by the model indicate that sustained activity in AIY can compensate for reduced AIB responses leading to robust odour approach. While it is likely that sexual conditioning may create many other changes within the circuit for odour processing in addition to the ones identified in this work, our data provides insight into how parallel memories of opposite valence can be resolved into a behavioural decision.

## Discussion

### Specificity of information coding by neuropeptide signalling

Here we have identified the mechanisms by which valence is flexibly modified by experience and neuropeptide signalling. We found that the neuropeptide PDF-1 can encode both positive and negative valence depending on its neuron source of release and the target neurons on which it acts. PDF-1 release from the interneurons MCM and AVB provides information about mate experience which acts as a teaching signal for reward. This teaching signal is incorporated into the circuit for odour processing by acting on the interneurons RIM, RIA and modulating the Ca^2+^ responses of the AIY axon upon odour stimulation. It is important to note that while AVB, a source of PDF-1, becomes activated during conditioning, the consequences of PDF-1 neuromodulation on AIY are observed during retrieval. This underscores the characteristic delay and slow kinetics with which neuropeptide signaling affects neural circuit function and behaviour. Modulation of this circuit by PDF-1 leads to odour approach. Odour avoidance on the other hand, requires the release of PDF-1 or PDF-2 from neurons other than AVB and MCM and modulation of interneurons other than RIM, RIA and AIY. More detailed mapping of the circuit for avoidance awaits further study. The orchestration of valence assignment by PDF-1 through the action on segregated circuits is reminiscent of the function of dopamine in the mushroom body of insects ^43, 44^, and in the ventral tegmental area of mammals ^45, 46^. In the amygdala as well, neuropeptides like neurotensin and cholecystokinin act on distinct circuits with segregated inputs and outputs to drive approach or avoidance ^47 48^. Therefore, the encoding of opposite valence by a single neuromodulator acting with circuit specificity appears to be a universal property of nervous systems. In mammals, the neuropeptide VIP, which belongs to the same family as PDF-1 ^49, 50^, is expressed in inhibitory interneurons with specific circuit topology ^51^. In cortex, amygdala and thalamo-cortical projections, VIP-expressing neurons become activated by behaviourally relevant salient stimuli to allow associative learning, long-term potentiation and memory retrieval ^52–54^. Although these functions have been previously assigned to the GABAergic neurotransmission of VIP-expressing neurons, in light of our results it will be important to investigate the role that neuromodulation through VIP plays in learning.

### The mechanisms underlying sex-specific learning

As previously shown by Sakai *et al*., we find that mate-experience is only rewarding for males since odour preferences can only be sexually conditioned in males. Perhaps a little surprising, this sexual dimorphism arises through activation of, and neuropeptide release from, a sex-shared neuron (AVB) and modulation of sex-shared neurons (RIM, RIA and AIY). Sex-specific activation of AVB is due to male-specific input connectivity from mate-sensing circuits. Although the male-specific MCM interneurons also receive input connectivity from mate-sensing circuits and are a required source of PDF-1, we did not observe changes in intracellular Ca^2+^ during conditioning. This suggests that other mechanisms may underlie their activation. A similar observation has been made for ASI neurons, which despite expressing receptors for pheromones and their function being required for pheromone-induced developmental and behavioural processes, do not display changes in intracellular Ca^2+^ levels upon pheromone exposure ^55–57^. Alternative activation mechanisms may include changes in gene expression ^58^ or membrane excitability ^59^. Nonetheless we cannot rule out the possibility that our experimental system may not have enough sensitivity to detect all meaningful Ca^2+^ changes. It will be important for future studies to determine whether there are sex differences in the ability of sex-shared neurons to respond to neuropeptide modulation to switch valence. Although sexually dimorphic innate behaviour has been extensively investigated, this is the first time that a multilevel (i.e. molecular, cellular, circuit and behavioural) analysis of sex differences in learning has been performed. We have been able to link the release of a neuropeptide from a particular class of neurons which are activated by male-specific input, to changes in neuronal activity and a switch in behaviour that underlies sex differences in learning.

### Reward and punishment create two memories that co-exist and compete for behavioural expression

Our results reveal that pairing a stimulus with both a punishment and a reward creates two parallel memories, one aversive and one rewarding, which compete for behavioural expression. The aversive memory, which arises from the association of an odour with the punishment of starvation and leads to odour avoidance, leaves a signature trace in the circuit characterised by changes in neural activity and trans-location of cellular components some of which have previously been shown to underlie avoidance ^39, 40^. Specifically, we observed lack of adaptation in RIA, a reduction in the number of AIB and RIM neurons that responded to odour removal, and nuclear translocation of PKG/EGL-4 in the odour-sensing neuron AWC. This signature, which is absent in naïve animals, remains present in animals that have been conditioned to associate the odour with both a punishment and the reward of mating partners (sexually conditioned animals) despite their behaviour being approach. In addition, in sexually conditioned animals, we observed a signature of reward characterised by perdurance in AIY axonal activity and that is dependent on PDF-1 neuromodulation. This, and possibly other, PDF-1-dependent traces are important to override the behavioural expression of the aversive memory, since removal of PDF-1 results in sexually conditioned animals that display no preference for or even avoidance of the odour. Although we have been able to identify signatures of both memories, there are likely many other neuronal changes occurring during conditioning which we have not extracted with our analysis and may account for the full behavioral expression of approach or avoidance. The idea that individual memories with opposite valence are simultaneously stored in different neurons to affect behaviour has been shown in the context of extinction learning ^60^ and spaced training ^61^. However, the synchronous association of opposite outcomes investigated here has only been studied at the level of behaviour ^62–65^. We have been able to show the specific changes that each experience produces in the brain and how these changes may regulate behaviour.

### Distinct neural representations of positive valence according to task functionality

Approach and avoidance are behavioural readouts of positive and negative valence, respectively. Research on how circuits encode valence has put forward two main functional types: those that act as modules and those that act as modes ^66^. Modules are those in which approach and avoidance are encoded by the activity of distinct and segregated circuits. Examples of modules have been identified in the amygdala and the ventral tegmental area of rodents ^47, 48, 67^ and the fly mushroom body ^68^. Modes are those in which avoidance or approach are encoded by the activation or inhibition of a neuron (or circuit, brain region) and examples have been found in the rodent dorsal raphe nucleus ^67^, *Lymnea* ^69^ and *C. elegans* ^25, 70^. Here, we do not find evidence of either modules or modes. We find no correlation between the activity of any particular neuron and the behavioural response to the odour. While it may be possible that we have not yet identified the neurons that encode odour valence, we think that this is unlikely. We have imaged the main interneurons of the circuit for odour processing and two of them, AIY and RIA, have previously been shown to encode experience-dependent changes in valence for CO_2_ and pathogenic bacteria ^25, 26^. Instead, what our results reveal is that positive valence can be flexibly represented according to functionality or predictions associated with a stimulus. While both naïve and sexually conditioned animals approach the odour benzaldehyde, odour sensation elicits very different neural activity which seems to reflect the different predictive value of the odour. Benzaldehyde, which is a bacteria metabolite, innately predicts food for naïve animals. In sexually conditioned animals in contrast, benzaldehyde predicts mates, and this is due to top-down modulation from downstream interneurons (AVB and MCM) to first layer interneurons (AIY). A similar process has been proposed for how sensory representation in the cortex changes according to task-specific brain states ^71^. Here we demonstrate that such flexible representation of sensory information can support learning and flexible behaviour.

## Methods

### *C. elegans* strains

Nematode culture and genetics were performed following standard procedures^72, 73^. A complete list of the strains used can be found in Table 2. All strains were in a *him-5* (e1490) background to facilitate male analysis. All assays were performed at 20ºC.

### Odour conditioning assays

Males and hermaphrodites were picked as L4s the night before the assay and transferred to single-sex plates with food. Animals were recovered from the food plates with wash buffer (1 mM CaCl_2_, 1 mM MgSO_4_, and 5 mM pH 6.0 potassium phosphate) and centrifuged at 1,700 r.p.m. for 3 min. A total of four washes were performed to remove the food before placing the animals on the conditioning plates.

Odour conditioning assay plates were made with 2% agar, 5 mM potassium phosphate (pH 6.0), 1 mM CaCl_2_ and 1 mM MgSO_4_. Mock conditioning plates were 5 cm in diameter (4 ml of agar), whereas aversive and sexual conditioning plates were 3 cm in diameter (2 ml of agar). Between 20-30 males were placed on each mock and aversive conditioning plate and exposed to either 100% ethanol or 15% benzaldehyde, respectively. For sexual conditioning, a maximum of 30 males and 200 hermaphrodites were used and animals were exposed to 15% benzaldehyde. Genotypes that were being compared against one another were sexually conditioned together in the same plate, tested and scored blindly and then identified through genotyping or by fluorescent markers. When different genotypes were conditioned together equal amounts of each genotype were used until a total of 30 males were added to the sexual conditioning plate. Controls in which animals were exposed to the odour on food were also performed for some experiments. In all cases animals were conditioned for 3 h.

Immediately after conditioning, animals were tested individually in 5 cm plates with a 1% benzaldehyde gradient by placing them in the centre of the assay plate, 2 cm away from the odour source (Fig. 1a). After 20 min, the tracks left by the animal were visualised and the chemotaxis score was calculated as previously described ^15^ by the sum of the scores of the regions by which the animal had travelled (Fig. 1a). Scoring of all sexual conditioning experiments were blind to the manipulation. All behavioural experiments were performed using N2 hermaphrodites. For the calcium imaging experiments *unc-51 (e369*) hermaphrodites were used for sexual conditioning.

### Plasmid constructs and transgenic strains

All constructs were built using Gibson Assembly ^74, 75^. Primers were designed using the NEBuilder assembly online tool (https://nebuilder.neb.com). The 3’ UTR of the *unc-54* gene was used for all constructs. Two different promoters were used to express *pdfr-1* in different cells. A promoter containing a 3kb region upstream of the translational *pdfr-1* start site (proximal promoter) drove *pdfr-1* isoforms b and d in neurons and body wall muscle ^21^ (Fig. 2b). A second construct drove a floxed isoform d in reverse orientation under a 5.1 kb sequence located 12.1Kb upstream of the translational *pdfr-1* start site (distal promoter) ^22^ (Fig. 2b). This construct was used to express *pdfr-1* cDNA isoform d in specific neurons through intersectional expression of Cre recombinase (Fig. 2b) A complete list of primers and plasmids used in this study can be found in Table 3 and Table 4, respectively. Plasmid constructs were injected in *C. elegans* following standard procedures^76, 77^. The specific co-injection marker used as well as the concentration at which each array was injected can be found in Table 2.

### CaMPARI measurements

*lite-1(ce314)*X; *him-5(e1490)*V; *oleEx58[pdf-1::Campari (25 ng/µl) + cc::GFP (30ng/µl)]* males were conditioned with 15% benzaldehyde in 3 cm plates with no food and with or without *unc-51* hermaphrodites for sexual and aversive conditioning, respectively. After 3 hrs, CaMPARI was photoconverted by exposing the plates to 405nm blue light (DAPI filter) for 2 minutes at 50% LED intensity under an inverted Zeiss Axiovert microscope and a 10x objective. Images of the MCM and AVB neurons were taken by mounting worms in 10% agarose pads with undiluted microbeads^78^. Classically used anesthetics such as NaN_3_ or levamisole could not be used because they interfere with the fluorescence signal of CaMPARI. Photos were taken with a Zeiss Axio Imager 2 microscope using a 470nm LED source at 25% intensity and EX470/20 FT493 EM505-503 filter for the green channel and GYR LED source at 50% intensity and BP546/12 FT580 LP590 filter for the red channel and at 125x magnification. LED intensity and exposure time were kept constant for AVB and MCM neurons across the sample population but longer exposure was needed for the MCM neurons. Image analysis was performed using Fiji. Before image processing, background intensity was subtracted from both channels. The average fluorescence intensity for each channel was measured by manually drawing a polygon containing the region of interest (ROI) on the z-plane at which the neuron analysed provided the brightest signal. The ratio between the average fluorescence intensity for green and red channels was calculated for each ROI.

### Calcium imaging

Animals were first subjected to one of the three conditioning protocols in the same way as for behavioural analysis. Immediately after conditioning, animals were injected into a nose trap microfluidic device ^34^ using wash buffer 1 mM CaCl_2_, 1 mM MgSO_4_, and 5 mM pH 6.0 potassium phosphate and 0.5 mM tetramisole to reduce movement of the animals while imaging. Animals were exposed to alternative periods of 10^−4^ benzaldehyde (diluted in wash buffer, no fluorescein) or wash buffer with 0.003 mg/ml fluorescein. Flow control buffer contained 0.03 mg/ml fluorescein. This allowed visualisation of flow with fluorescence, and control checks of adequate flow were done between every recording. Imaging was performed in an inverted Zeiss Axiovert microscope with a 460/565nm LED. Emission filters ET515/30M and ET641/75 and dichroic T565lprx-UF2 were placed in the cube of a Cairn OptoSplit II attached between the microscope and an ORCA-Flash four camera (Hamamatsu). Acquisition was performed at 10 fps with a 63X objective (Zeiss plan-apochromat, numerical aperture 1.4) and 1.5X optovar. To avoid photobleaching pulse illumination was used. The cell body was imaged for AIB neurons, whereas the projection was imaged for RIM, AIY (specifically zone 2), and RIA (loop domain, ventral nerve ring domain and dorsal nerve ring domain).

### Microfluidic chip fabrication

Male-adapted PDMS olfactory chips ^35^ were made and plasma-bonded to a glass coverslip as previously described ^35, 79^. After plasma bonding, chips were placed on a hot plate at 80 ºC for one hour to improve bonding efficiency.

### Calcium imaging analysis

A moving region of interest in both channels was identified and mean fluorescent ratios (GFP/RFP) were calculated with custom-made Matlab scripts by the de Bono lab ^80^. All further analysis was performed using our own custom-made Matlab scripts. For RIA neurons, where photobleaching was evident, bleach-correction was applied independently to each channel by fitting an exponential decay curve to the raw fluorescence signal and dividing the signal by its fitted exponential ^81^. Ratios were smoothened using a 5-frame rolling median. Baseline-adjusted ratios were calculated as (R-R0)/R0, where R0 is the average ratio during the baseline period (10s prior to odour presentation for AIY and RIA, and 10s prior to odour removal for RIM and AIB). Normalised ratios were calculated as (R-Rmin)/Rmax, where Rmin and Rmax are the average of the smallest and largest 5% of values in that trace, respectively. For AIB and RIM, normalised ratios were calculated in order to classify neurons as being in either their ON or OFF state (classed as ON if (R-Rmin)/Rmax > 0.5, as per ^24, 26^, and then baseline-adjusted ratios were plotted to show differences in magnitude of response across conditions.

### EGL-4 quantification

Animals were first subjected to one of the four conditioning protocols in the same way as for behavioural analysis. Immediately after conditioning, animals were split in two groups. One group was imaged for EGL-4 localization and the other group was tested for chemotaxis to benzaldehyde. Imaging of GFP-tagged EGL-4 was performed with a Zeiss Axio Imager 2 microscope, a Zeiss ECPLAN-NEOFLUAR 100x/1.3 oil immersion objective and TimeToLive 10_2016 Caenotec software. Animals were immobilised with 3μl of NaN_3_ at a concentration of 25mM. AWC neurons were identified by expression of *odr-1::dsRed*, morphology and position. Z-stack images were taken for each animal and analysed using ImageJ blindly to the conditioning that the worms had been subject to. Average green fluorescence intensity at nuclear and cytoplasmic locations was calculated by drawing a polygon containing each region of interest (ROI) on the middle section of the z-stack for each AWC neuron and the ratio was calculated. Animals with ratios ≥ 1,4 were classified as having EGL-4 nuclear translocation.

### Statistical analysis

For behavioural assays, because chemotaxis scores are discrete categorical data, ξ_2_ for trend analysis was used to compare the frequency distribution of values of different conditions. Bonferroni corrections were applied to account for multiple comparisons-derived false positives.

For CaMPARI photo conversion, a non-parametric two tailed Mann Whitney-U test was performed.

To compare EGL-4 nuclear translocation between different conditioning groups a ξ_2_ test with Bonferroni corrections was used to account for multiple comparisons derived false positives.

For GCaMP fluorescence analysis, non-parametric one-way ANOVA (Kruskal Wallis test) with Dunn’s multiple comparisons was performed because the data did not follow a normal distribution. The same test was used to compare the proportion of AIB and RIM neurons activated over time. χ^2^ test was used to compare the proportion of type 2 responses in AIY between conditions and genotypes. The type of test used for each dataset is specified in each figure legend.

### Finite difference model of worm behaviour

To investigate the worm’s behaviour an ordinary differential equation model was developed (Extended Data Fig. 4). This simulated the worm’s response to a concentration gradient by modelling a simple neural network. The network was comprised of the AWC sensory neurons and the AIB, AIA and AIY neurons. We adapted the modelling approach in ^82^. The sensory neuron AWC is comprised of two components which react to the concentration of odour on different timescales, a fast, *F*, and slow, *S*, component^82^. The input to the sensory neuron is given by

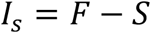

The fast component integrates over a sensory stimulus *C* with a characteristic rate *α* and decay rate *β*

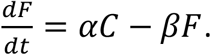

The slow component moves towards the fast component with a delay

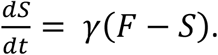

The intermediate neurons AIB, AIA, AIY were modelled according to the approach in ^82^. All neurons are modelled as leaky integrators:

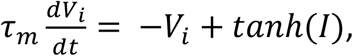

where *V*_*i*_ is the voltage or activity of neuron *i, τ*_*m*_ is a time constant, *V*_*0,i*_ is the resting potential and *I* is an input term. The input term is a sum over all inputs to a neuron *I* = ∑_*j*_ *w*_*ij*_ *V*_*j*_. The parameter values were *α* = 4s^−1^, *β* = 15s^−1^, *γ* = *2*s^−1^, *t*_*m*_ = 0.5s ^82^

AIB activation is known to correspond to reorientation behaviour in the worm and AIY activation to cause the worm to move forwards along its current trajectory. To include this in the model the worms make stochastic decisions in each modelled time step to either continue going straight forward or to reorient to a new, random orientation. The probability of reorientation is governed by the response neuron R which is influenced by the activation of AIB and AIY. At each time step the worm can choose to continue moving forward on its current direction or to reorient itself by choosing a new random direction uniformly from 0-360 degrees. This decision is made by the reorientation neuron R, the output of which is

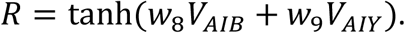

To make a reorientation decision a random number is drawn uniformly between −1 and 1. If this number is lower than *R* the worm will reorient, otherwise the worm will continue moving in its current direction.

Worm movement was simulated on a circular plate with radius 30 mm using the forward Euler method with a timestep *dt* = 0.005*s*. A worm starts at position (0,0) with an initial direction of motion, *θ*, drawn uniformly from 0-360 degrees. Worms move with a speed of *v* = 0.11*mms*^−1 82^, and at every timestep a reorientation decision is made. If the worm decides to reorient a new direction of motion is sampled from 0-360 degrees and the worm starts moving in this direction, otherwise the worm continues on its current direction of motion. The concentration of odour is given by a gaussian function with origin at (4.5, 0) and standard deviation of 4mm *C*(*x, y*) = 100 ∗ *N*(4.5, *0*, 4).

To compare the simulated worms to the real worms a fitness function (*F*) was constructed using summary statistics of the distribution of sectors visited by the simulated and real worms. The score of a worm is the sum of all sectors that it visited. The range of a worm is the difference between the highest and lowest sector visited. The fitness function of a set of weights is based on the mean, standard deviation and skew of the score and the mean, standard deviation and skew of the range of the worms in a simulated experiment compared to the real data.

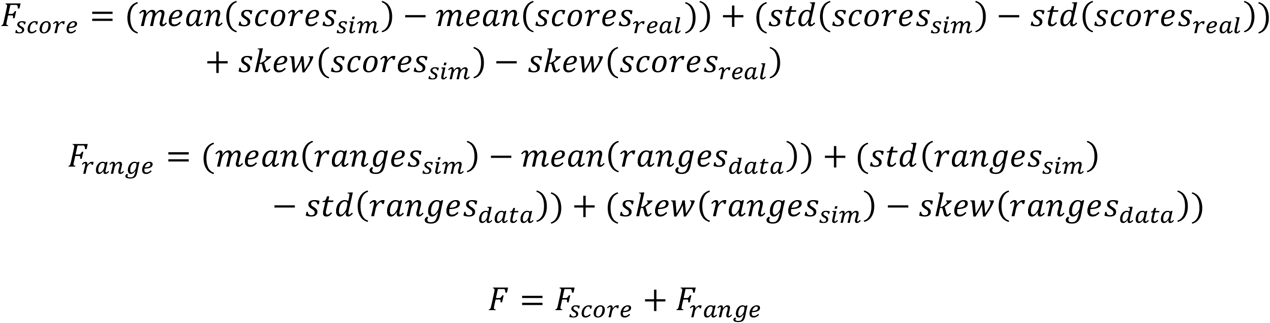

First data for 92 worms with no odour applied was used to find an effective sampling time for the worms. Simulated experiments of 92 worms were carried out for sampling time [0.06, 0.07, 0.08, 0.09, 0.10, 0.11, 0.12, 0.13, 0.14] seconds (Extended Data Fig. 4c). A sampling time of 0.10 was found to have the highest fitness and therefore was used as the sampling time for all subsequent simulations.

The method to fit the weights of the neural network to different behaviours is shown in Extended Data Fig. 4d. The target behaviour is defined by 212 behavioural assay data points of mock conditioned worms. The evolutionary algorithm starts from a population of randomly initialised weights, however *w*_8_ was always set to 1 and *w*_9_ to −1. During each round of evolution the fitness of each set of weights in the population was calculated by carrying out a simulated experiment and calculating the fitness of the weights. The population was then ranked according to fitness. The top 40% of the population is left unchanged to go onto the next generation. The next 40% of the population is replaced by the top 40% after each weight has been perturbed by small random numbers. The final 20% of the population is replaced by new randomly sampled weights. The output of the evolutionary algorithm is the final population of weights. For all simulations the population size was 100 and it was evolved for 100 generations.

## Acknowledgements

We thank the following labs and individuals for strains and reagents: Bargmann (The Rockefeller University), L’Étoile (University of California San Francisco), Carrera (Universidad de la Republica), Srinivasan (Worcester Polytechnic Institute), García (Texas A&M University), Hobert (Columbia University), Ch’ng (Kings College London), Poole (UCL), Busch (Health and Medical University, Berlin), Temmerman (KU Leuven), Barkoulas (Imperial College), Iino (University of Tokyo), Sengupta (Brandeis University), Rosie Truman (UCL), Evie Goss-Sampson and Michele Sammut (UCL). Additional strains were obtained from the Caenorhabditis Genetic Center (University of Minnesota) which is supported by the National Institutes of Health - Office of Research Infrastructure Programs (P40 OD010440). We also thank Chintan Trivedi (UCL) for help setting up the odour stimulation system. We are grateful to H. Buelow and our colleagues at UCL J. Rihel, M. Amoyel and V. Fernandes for insightful comments on the manuscript. This work was supported by a Newton Fellowship from the Royal Society to LMG (NF160914), a Wellcome Trust PhD Studentship to SCF (175261) and a Leverhulme Trust project grant RPG-2018–287 to AB.

## Author contributions

A.B. conceived the project. A.B. and L.M.G. designed experiments, interpreted data and co-wrote the manuscript. L.M.G. built the strains, performed the behavioural experiments, and analysed data. L.M.G. and S.C.F. performed the neuronal recordings, interpreted and analysed the data. S.C.F. developed custom MatLab scripts to analyse the calcium imaging data. E.C., L.L., S.B.L. and B.G.M.O. contributed to imaging and behavioural experiments. N.T. and C.B. developed the computational model of odourtaxis.

## References

1. Dal Bello, M., Perez-Escudero, A., Schroeder, F.C. & Gore, J. Inversion of pheromone preference optimizes foraging in C. elegans. Elife 10 (2021).

2. Hughes, D.P. & Libersat, F. Parasite manipulation of host behavior. Curr Biol 29, R45–R47 (2019).

3. Keleman, K., et al. Dopamine neurons modulate pheromone responses in Drosophila courtship learning. Nature 489, 145–149 (2012).

4. Zhang, Y., Lu, H. & Bargmann, C.I. Pathogenic bacteria induce aversive olfactory learning in Caenorhabditis elegans. Nature 438, 179–184 (2005).

5. Ha, H.I., et al. Functional organization of a neural network for aversive olfactory learning in Caenorhabditis elegans. Neuron 68, 1173–1186 (2010).

6. Li, Q. & Liberles, S.D. Aversion and attraction through olfaction. Curr Biol 25, R120–R129 (2015).

7. Tye, K.M. Neural circuit motifs in valence processing. Neuron 100, 436–452 (2018).

8. Wagle, M., et al. Brain-wide perception of the emotional valence of light is regulated by distinct hypothalamic neurons. Mol Psychiatry 27, 3777–3793 (2022).

9. Carew, T.J., Walters, E.T. & Kandel, E.R. Classical conditioning in a simple withdrawal reflex in Aplysia californica. J Neurosci 1, 1426–1437 (1981).

10. Ward, S. Chemotaxis by the nematode Caenorhabditis elegans: identification of attractants and analysis of the response by use of mutants. Proc Natl Acad Sci U S A 70, 817–821 (1973).

11. Bargmann, C.I., Hartwieg, E. & Horvitz, H.R. Odorant-selective genes and neurons mediate olfaction in C. elegans. Cell 74, 515–527 (1993).

12. Tomioka, M., et al. The insulin/PI 3-kinase pathway regulates salt chemotaxis learning in Caenorhabditis elegans. Neuron 51, 613–625 (2006).

13. Nuttley, W.M., Atkinson-Leadbeater, K.P. & Van Der Kooy, D. Serotonin mediates food-odor associative learning in the nematode Caenorhabditis elegans. Proc Natl Acad Sci U S A 99, 12449–12454 (2002).

14. Lin, C.H., et al. Insulin signaling plays a dual role in Caenorhabditis elegans memory acquisition and memory retrieval. J Neurosci 30, 8001–8011 (2010).

15. Sakai, N., et al. A sexually conditioned switch of chemosensory behavior in C. elegans. PLoS One 8, e68676 (2013).

16. Sammut, M., et al. Glia-derived neurons are required for sex-specific learning in C. elegans. Nature 526, 385–390 (2015).

17. Barrios, A., Nurrish, S. & Emmons, S.W. Sensory regulation of C. elegans male mate-searching behavior. Curr Biol 18, 1865–1871 (2008).

18. Lipton, J., Kleemann, G., Ghosh, R., Lints, R. & Emmons, S.W. Mate searching in Caenorhabditis elegans: a genetic model for sex drive in a simple invertebrate. J Neurosci 24, 7427–7434 (2004).

19. Janssen, T., et al. Functional characterization of three G protein-coupled receptors for pigment dispersing factors in Caenorhabditis elegans. J Biol Chem 283, 15241–15249 (2008).

20. Janssen, T., et al. Discovery and characterization of a conserved pigment dispersing factor-like neuropeptide pathway in Caenorhabditis elegans. J Neurochem 111, 228–241 (2009).

21. Barrios, A., Ghosh, R., Fang, C., Emmons, S.W. & Barr, M.M. PDF-1 neuropeptide signaling modulates a neural circuit for mate-searching behavior in C. elegans. Nat Neurosci 15, 1675–1682 (2012).

22. Flavell, S.W., et al. Serotonin and the neuropeptide PDF initiate and extend opposing behavioral states in C. elegans. Cell 154, 1023–1035 (2013).

23. Chalasani, S.H., et al. Dissecting a circuit for olfactory behaviour in Caenorhabditis elegans. Nature 450, 63–70 (2007).

24. Gordus, A., Pokala, N., Levy, S., Flavell, S.W. & Bargmann, C.I. Feedback from network states generates variability in a probabilistic olfactory circuit. Cell 161, 215–227 (2015).

25. Guillermin, M.L., Carrillo, M.A. & Hallem, E.A. A single set of interneurons drives opposite behaviors in C. elegans. Curr Biol 27, 2630–2639 e2636 (2017).

26. Jin, X., Pokala, N. & Bargmann, C.I. Distinct circuits for the formation and retrieval of an imprinted olfactory memory. Cell 164, 632–643 (2016).

27. Brockie, P.J., Madsen, D.M., Zheng, Y., Mellem, J. & Maricq, A.V. Differential expression of glutamate receptor subunits in the nervous system of Caenorhabditis elegans and their regulation by the homeodomain protein UNC-42. J Neurosci 21, 1510–1522 (2001).

28. Hobert, O., et al. Regulation of interneuron function in the C. elegans thermoregulatory pathway by the ttx-3 LIM homeobox gene. Neuron 19, 345–357 (1997).

29. Cook, S.J., et al. Whole-animal connectomes of both Caenorhabditis elegans sexes. Nature 571, 63–71 (2019).

30. Srinivasan, J., et al. A blend of small molecules regulates both mating and development in Caenorhabditis elegans. Nature 454, 1115–1118 (2008).

31. Liu, K.S. & Sternberg, P.W. Sensory regulation of male mating behavior in Caenorhabditis elegans. Neuron 14, 79–89 (1995).

32. Barr, M.M. & Sternberg, P.W. A polycystic kidney-disease gene homologue required for male mating behaviour in C. elegans. Nature 401, 386–389 (1999).

33. Fosque, B.F., et al. Neural circuits. Labeling of active neural circuits in vivo with designed calcium integrators. Science 347, 755–760 (2015).

34. Chronis, N., Zimmer, M. & Bargmann, C.I. Microfluidics for in vivo imaging of neuronal and behavioral activity in Caenorhabditis elegans. Nat Methods 4, 727–731 (2007).

35. Reilly, D.K., Lawler, D.E., Albrecht, D.R. & Srinivasan, J. Using an adapted microfluidic olfactory chip for the imaging of neuronal activity in response to pheromones in male C. elegans head neurons. J Vis Exp (2017).

36. Chalasani, S.H., et al. Neuropeptide feedback modifies odor-evoked dynamics in Caenorhabditis elegans olfactory neurons. Nat Neurosci 13, 615–621 (2010).

37. Liu, H., et al. Cholinergic sensorimotor integration regulates olfactory steering. Neuron 97, 390–405 e393 (2018).

38. Gray, J.M., Hill, J.J. & Bargmann, C.I. A circuit for navigation in Caenorhabditis elegans. Proc Natl Acad Sci U S A 102, 3184–3191 (2005).

39. Lee, J.I., et al. Nuclear entry of a cGMP-dependent kinase converts transient into long-lasting olfactory adaptation. Proc Natl Acad Sci U S A 107, 6016–6021 (2010).

40. Cho, C.E., Brueggemann, C., L’Etoile, N.D. & Bargmann, C.I. Parallel encoding of sensory history and behavioral preference during Caenorhabditis elegans olfactory learning. Elife 5 (2016).

41. Tsalik, E.L. & Hobert, O. Functional mapping of neurons that control locomotory behavior in Caenorhabditis elegans. J Neurobiol 56, 178–197 (2003).

42. Ben Arous, J., Laffont, S. & Chatenay, D. Molecular and sensory basis of a food related two-state behavior in C. elegans. PLoS One 4, e7584 (2009).

43. Cohn, R., Morantte, I. & Ruta, V. Coordinated and compartmentalized neuromodulation shapes sensory processing in Drosophila. Cell 163, 1742–1755 (2015).

44. Traniello, I.M., Chen, Z., Bagchi, V.A. & Robinson, G.E. Valence of social information is encoded in different subpopulations of mushroom body Kenyon cells in the honeybee brain. Proc Biol Sci 286, 20190901 (2019).

45. de Jong, J.W., et al. A neural circuit mechanism for encoding aversive stimuli in the mesolimbic dopamine system. Neuron 101, 133–151 e137 (2019).

46. McGovern, D.J., Polter, A.M. & Root, D.H. Neurochemical signaling of reward and aversion to ventral tegmental area glutamate neurons. J Neurosci 41, 5471–5486 (2021).

47. Li, H., et al. Neurotensin orchestrates valence assignment in the amygdala. Nature 608, 586–592 (2022).

48. Zhang, X., et al. Genetically identified amygdala-striatal circuits for valence-specific behaviors. Nat Neurosci 24, 1586–1600 (2021).

49. Frooninckx, L., et al. Neuropeptide GPCRs in C. elegans. Front Endocrinol (Lausanne) 3, 167 (2012).

50. Janssen, T., Lindemans, M., Meelkop, E., Temmerman, L. & Schoofs, L. Coevolution of neuropeptidergic signaling systems: from worm to man. Ann N Y Acad Sci 1200, 1–14 (2010).

51. Pi, H.J., et al. Cortical interneurons that specialize in disinhibitory control. Nature 503, 521–524 (2013).

52. Krabbe, S., et al. Adaptive disinhibitory gating by VIP interneurons permits associative learning. Nat Neurosci 22, 1834–1843 (2019).

53. Melzer, S., et al. Bombesin-like peptide recruits disinhibitory cortical circuits and enhances fear memories. Cell 184, 5622–5634 e5625 (2021).

54. Williams, L.E. & Holtmaat, A. Higher-order thalamocortical inputs gate synaptic long-term potentiation via disinhibition. Neuron 101, 91–102 e104 (2019).

55. Kim, K., et al. Two chemoreceptors mediate developmental effects of dauer pheromone in C. elegans. Science 326, 994–998 (2009).

56. Luo, J. & Portman, D.S. Sex-specific, pdfr-1-dependent modulation of pheromone avoidance by food abundance enables flexibility in C. elegans foraging behavior. Curr Biol 31, 4449–4461 e4444 (2021).

57. McGrath, P.T., et al. Parallel evolution of domesticated Caenorhabditis species targets pheromone receptor genes. Nature 477, 321–325 (2011).

58. Laurent, P., et al. Decoding a neural circuit controlling global animal state in C. elegans. Elife 4 (2015).

59. Stout, R.F., Jr., Verkhratsky, A. & Parpura, V. Caenorhabditis elegans glia modulate neuronal activity and behavior. Front Cell Neurosci 8, 67 (2014).

60. Felsenberg, J., et al. Integration of parallel opposing memories underlies memory extinction. Cell 175, 709–722 e715 (2018).

61. Jacob, P.F. & Waddell, S. Spaced training forms complementary long-term memories of opposite valence in Drosophila. Neuron 106, 977–991 e974 (2020).

62. Burgos-Robles, A., et al. Amygdala inputs to prefrontal cortex guide behavior amid conflicting cues of reward and punishment. Nat Neurosci 20, 824–835 (2017).

63. Aso, Y. & Rubin, G.M. Dopaminergic neurons write and update memories with cell-type-specific rules. Elife 5 (2016).

64. Das, G., et al. Drosophila learn opposing components of a compound food stimulus. Curr Biol 24, 1723–1730 (2014).

65. Stephenson-Jones, M., et al. Opposing contributions of GABAergic and Glutamatergic ventral pallidal neurons to motivational behaviors. Neuron 105, 921–933 e925 (2020).

66. Berridge, K.C. Affective valence in the brain: modules or modes? Nat Rev Neurosci 20, 225–234 (2019).

67. Li, Y., et al. Rostral and Caudal Ventral Tegmental Area GABAergic inputs to different Dorsal Raphe neurons participate in opioid dependence. Neuron 101, 748–761 e745 (2019).

68. Waddell, S. Reinforcement signalling in Drosophila; dopamine does it all after all. Curr Opin Neurobiol 23, 324–329 (2013).

69. Pirger, Z., et al. Interneuronal mechanisms for learning-induced switch in a sensory response that anticipates changes in behavioral outcomes. Curr Biol 31, 1754–1761 e1753 (2021).

70. Fagan, K.A., et al. A single-neuron chemosensory switch determines the valence of a sexually dimorphic sensory behavior. Curr Biol 28, 902–914 e905 (2018).

71. Zhang, N. & Xu, N.L. Reshaping sensory representations by task-specific brain states: Toward cortical circuit mechanisms. Curr Opin Neurobiol 77, 102628 (2022).

72. Brenner, S. The genetics of Caenorhabditis elegans. Genetics 77, 71–94 (1974).

73. Stiernagle, T. Maintenance of C. elegans. WormBook, 1–11 (2006).

74. Gibson, D.G., et al. Enzymatic assembly of DNA molecules up to several hundred kilobases. Nat Methods 6, 343–345 (2009).

75. Gibson, D.G., et al. Creation of a bacterial cell controlled by a chemically synthesized genome. Science 329, 52–56 (2010).

76. Mello, C.C., Kramer, J.M., Stinchcomb, D. & Ambros, V. Efficient gene transfer in C. elegans: extrachromosomal maintenance and integration of transforming sequences. EMBO J 10, 3959–3970 (1991).

77. Berkowitz, L.A., Knight, A.L., Caldwell, G.A. & Caldwell, K.A. Generation of stable transgenic C. elegans using microinjection. J Vis Exp (2008).

78. Kim, E., Sun, L., Gabel, C.V. & Fang-Yen, C. Long-term imaging of Caenorhabditis elegans using nanoparticle-mediated immobilization. PLoS One 8, e53419 (2013).

79. Lagoy, R.C. & Albrecht, D.R. Microfluidic devices for behavioral analysis, microscopy, and neuronal imaging in Caenorhabditis elegans. Methods Mol Biol 1327, 159–179 (2015).

80. Busch, K.E., et al. Tonic signaling from O(2) sensors sets neural circuit activity and behavioral state. Nat Neurosci 15, 581–591 (2012).

81. Kaplan, H.S., Salazar Thula, O., Khoss, N. & Zimmer, M. Nested neuronal dynamics orchestrate a behavioral hierarchy across timescales. Neuron 105, 562–576 e569 (2020).

82. Ghosh, D.D., et al. Neural architecture of hunger-dependent multisensory decision making in C. elegans. Neuron 92, 1049–1062 (2016).

