## Supplemental material for "Conflict during learning reconfigures the neural representation of positive valence and approach behaviour"

**Extended Data Fig. 1: The MCMs and AVBs are not required sources of PDF-1 for aversive learning.** Chemotaxis score to benzaldehyde of *pdf-1(tm1996); pdf-2 (tm4393)* double mutant males after conditioning. The presence or absence of the excisable *pdf-1* transgene as a single copy insertion (syb1393) is indicated. The promotors used in the Cre transgenes are also indicated. Each empty circle within a violin plot represents the chemotaxis score of an individual animal. The mode is marked with a white cross. $\chi^2$ for trend analysis and Bonferroni correction was used to compare the frequency distribution of values of each genotype between conditions (black asterisks) and for a condition between genotypes (green asterisks). *** $p< 0.0001$; n. s. no statistically significant difference $p\geq0.05$. n, number of animals. Different genotypes were assayed in different days but always with a positive and a negative control. Data of the control groups are pulled together. At least three independent experiments were done for each genotype.

**Extended Data Fig. 2: AIB and RIM neurons alternate between high and low activity states.** **a** and **b,** Probability distribution of normalised $((R-R_{min})/R_{max})$ GCaMP6f/RFP fluorescence ratios showing bimodal distribution of activity in AIB (a) and RIM (b). **c** and **d,** Proportion of AIB (c) and RIM (d) neurons that became activated at each time point during rest, in the absence of odour exposure in mock (grey lines), aversively (green lines) and sexually conditioned (red lines) males. Non-parametric ANOVA was used to compared conditions. 75 neurons were analysed for mock, 42 for aversive, and 48 for sexual conditioning for AIB. 36 neurons were analysed for mock, 29 for aversive, and 40 for sexual conditioning for RIM. n. s. no statistically significant difference $p\geq0.05$.

**Extended Data Fig. 3: Sexual conditioning changes the responses of AIY to odour in a PDF-1-dependent manner**. **a,** Distribution of time to maximum response of normalised GCaMP6f/RFP fluorescence ratios $(R-R_{min})/R_{max})$ in the axon of AIY interneurons upon odour exposure after conditioning. Each dot represents an individual neuron and these are colour-coded according to their maximum activation $(R_{max})$. **b,** Proportion of type 2 responses in the axon of AIY interneurons of *pdf-1 (tm1996)* mutants. $\chi^2$ test was used to compare groups. n. s. no statistically significant difference $p\geq0.05$. 35 neurons were analysed for mock, 36 for aversive, and 50 for sexual conditioning. **c,** Average normalised traces of GCaMP6f/RFP fluorescence ratios $((R-R_{min})/R_{max})$ in the axons of AIY neurons of *pdf-1 (tm1996)* mutants after mock and aversive conditioning. Type 1 and type 2 responses have been plotted separately for each condition and their frequency indicated.

**Extended Data Fig. 4: A finite difference model of worm behaviour. a,** The neural network model. The sensory neuron AWC senses the odorant benzaldehyde and is activated by a down step in concentration. First layer interneurons AIB, AIA and AIY respond to the activation of AWC where the response is dependent upon the weights connected to these neurons. AIB and AIY then signal the response neuron R which stochastically makes the decision to reorient or continue moving forward. **b,** The simulated experimental setup. Worm motion is confined within a circular plate with radius of 3 mm (black circle). A worm is scored by summing the score of the sectors it passes through, sectors are separated by gray lines. When odour is applied this is simulated with a gaussian concentration field originating at location (4.5,0) with a standard deviation of 4. **c,** The results of the sample time fitting. **d,** Outline of the evolutionary algorithm used to find weights which match the behaviour of the worms.

**Extended Data Fig. 5: Simulated neuronal responses.** Neural voltage in AWC, AIB and AIY in each of the 100 simulated populations of worms. Simulated worms with sustained activity in AIY and reduced activity in AIB are labelled in green.

**Extended Data Fig. 6: Simulated odourtaxis behaviour.** Violin plots showing the simulated odourtaxis behaviour in each of the 100 simulated populations of worms (orange) overlaid onto the observed data of mock conditioned animals (blue). Simulations corresponding to sustained AIY activity and reduced AIB activity are labelled in green. Bars within violin plots represent the 10 and 90 percentile of each distribution.

**Extended Data Fig. 7: Correlation between observed and simulated chemotaxis scores.** QQ plots comparing the probability distribution of chemotaxis scores in the simulated and observed worm population. Simulations corresponding to sustained AIY activity and reduced AIB activity are labelled in green.

### Extended Data Fig.1

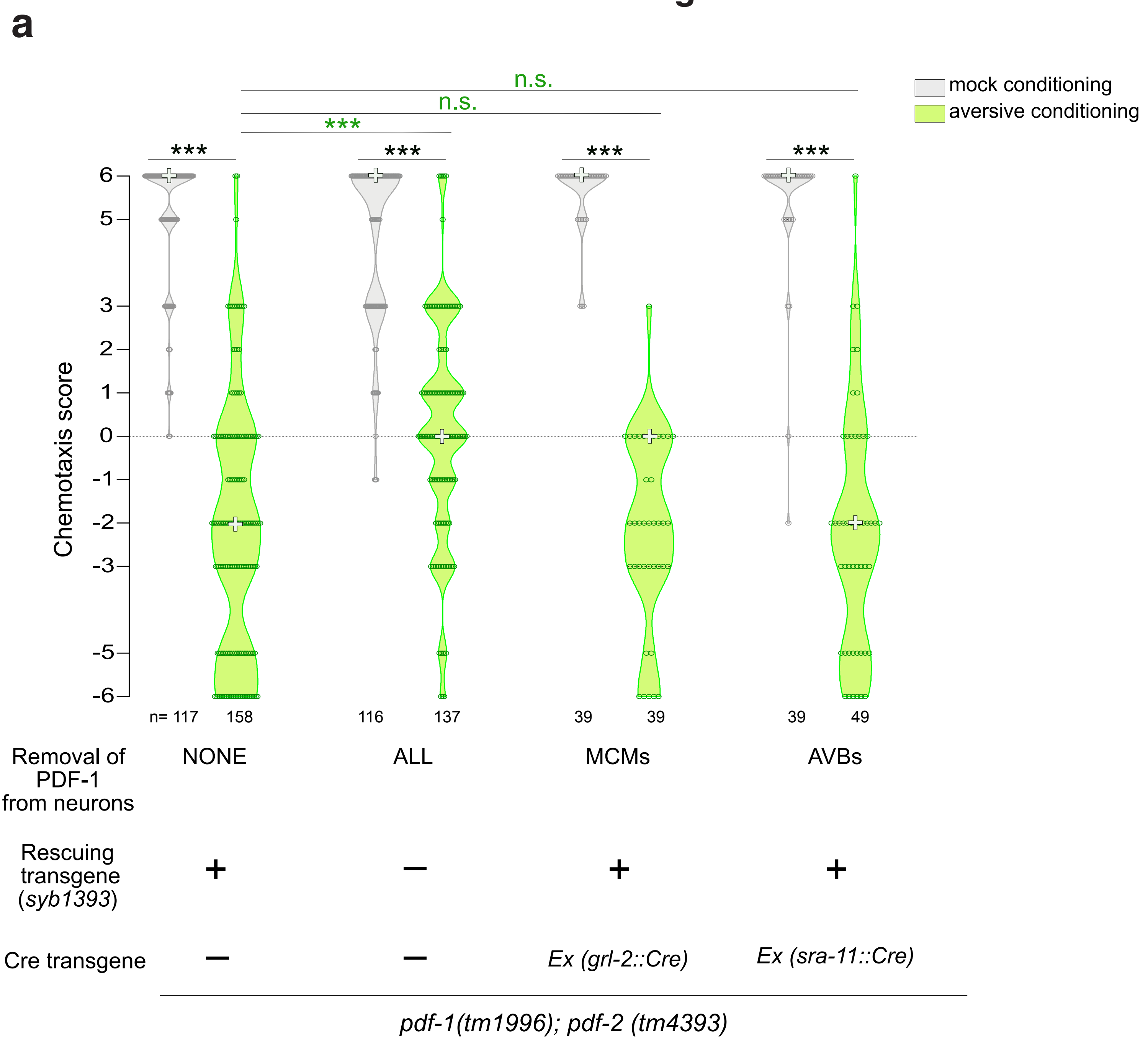

### Extended Data Fig.2

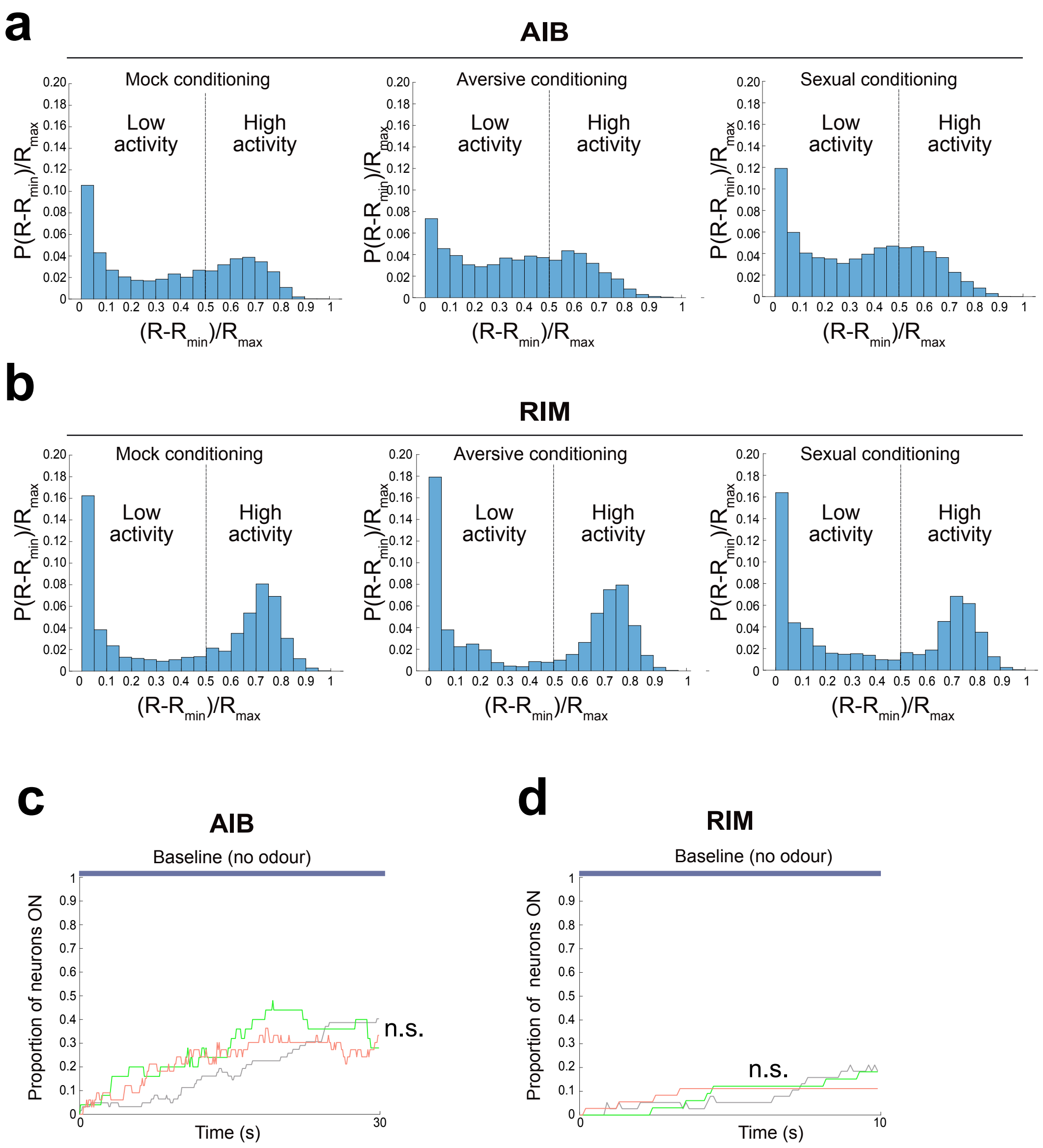

### Extended Data Fig.3

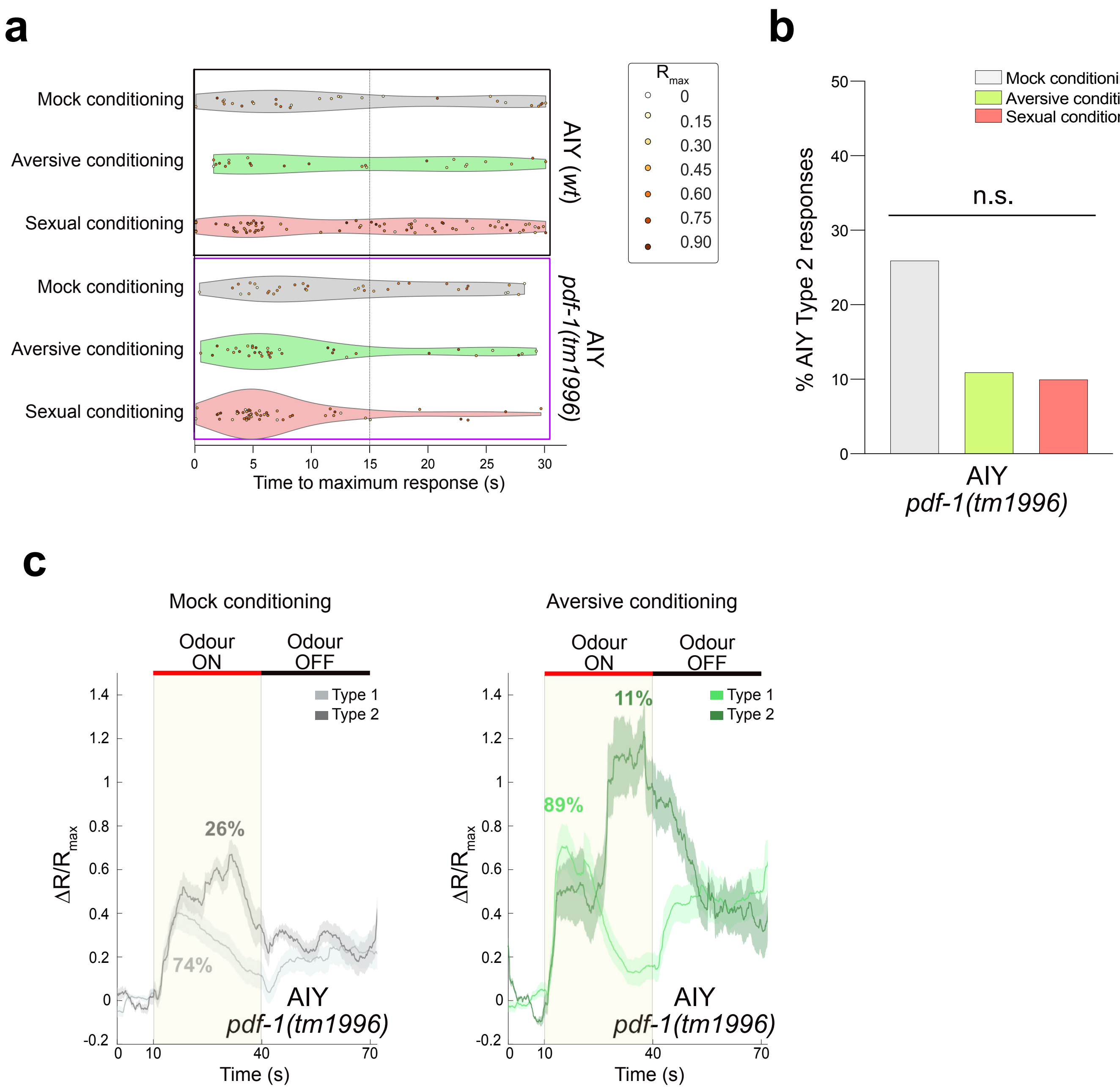

**Extended Data Fig. 4**

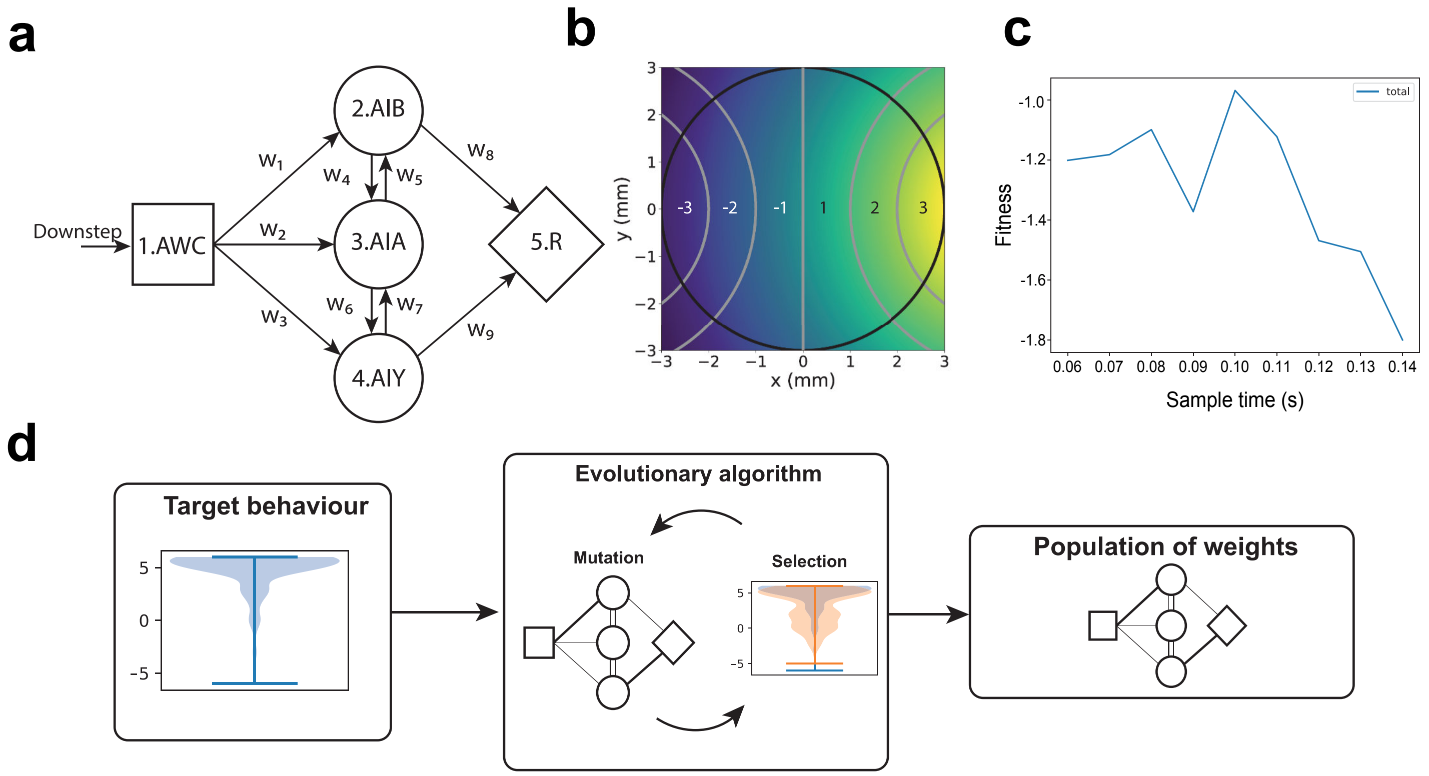

### Extended Data Fig. 5

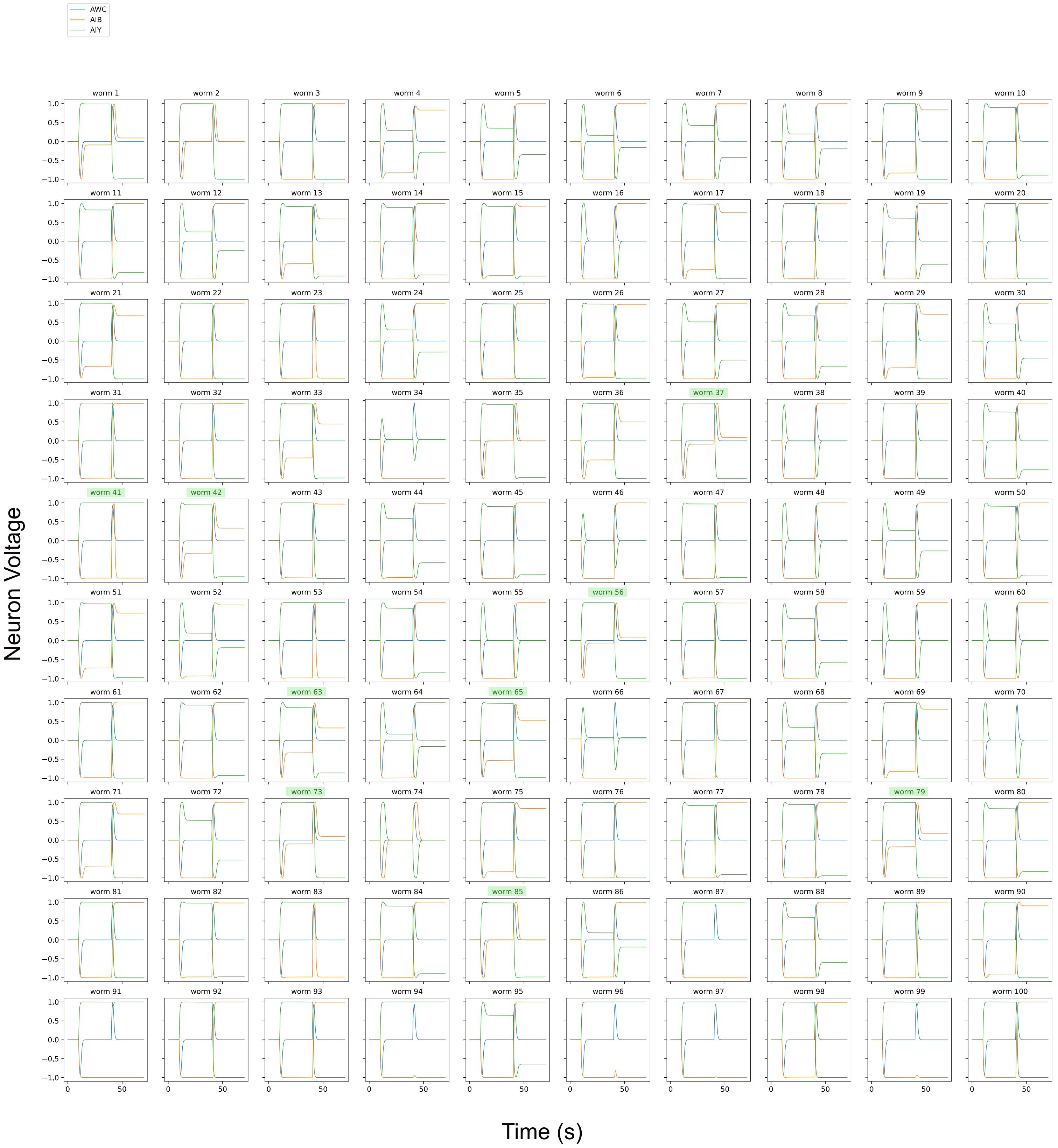

### Extended Data Fig. 6

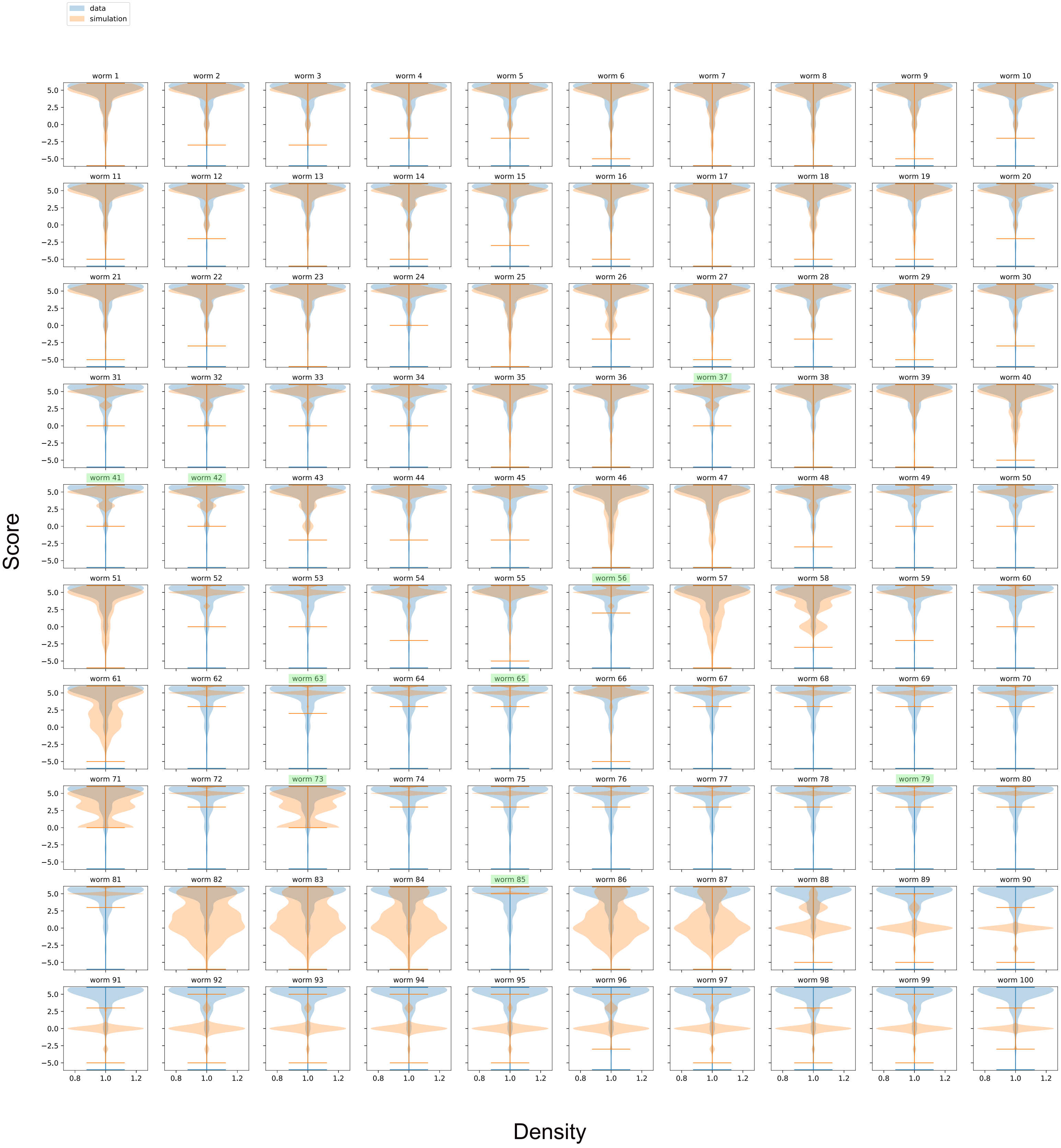

### Extended Data Fig. 7

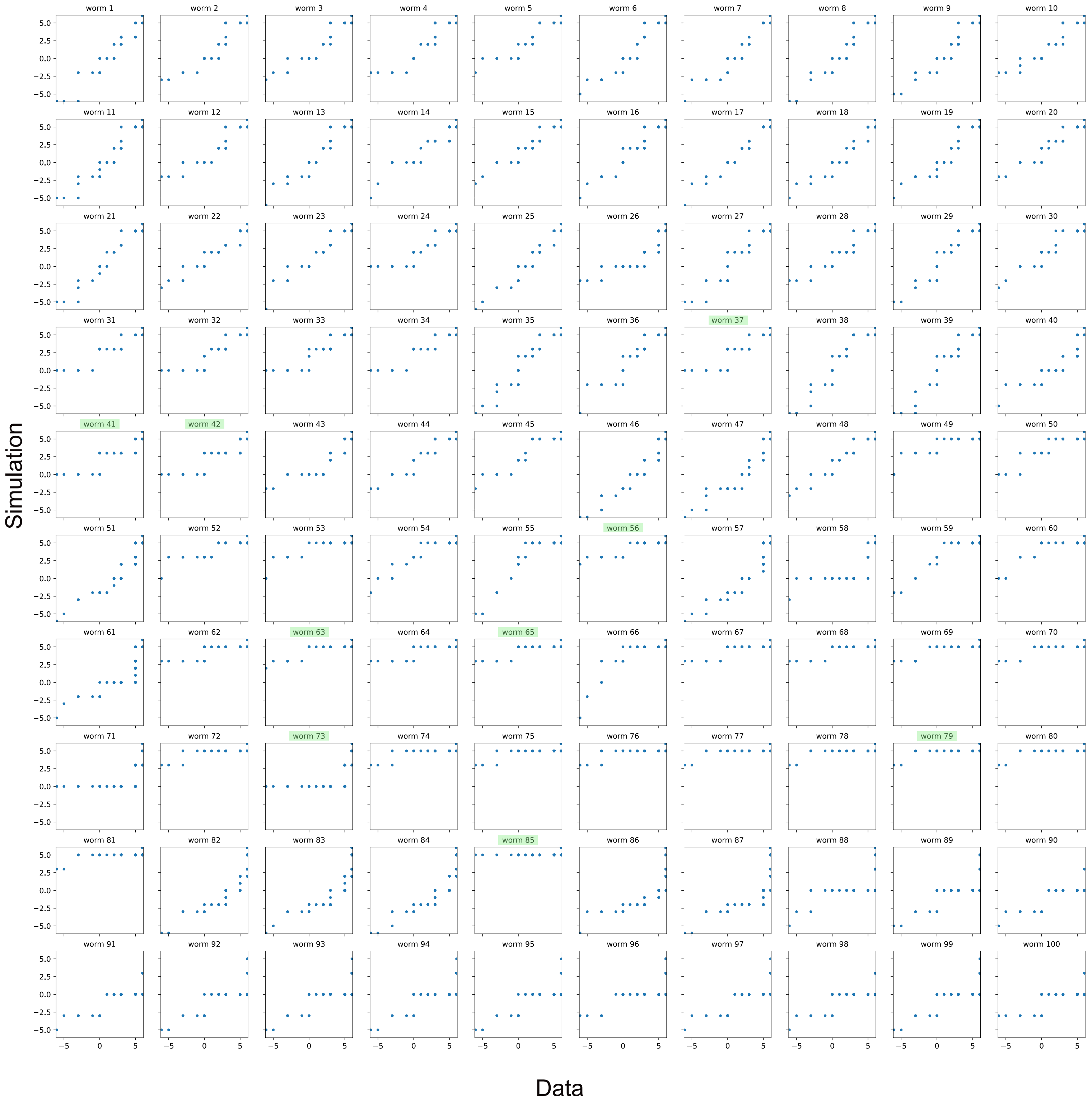

| Combination | | | Average chemotaxis score to 1% BZ | |
|---|---|---|---|---|
| Volume μl | Concentration (% in ethanol) | Time (h) | Mock conditioned | Aversively conditioned |
| 2.0 | 100 | 1 | 6.0 | Paralysed |
| 1.0 | 100 | 1 | 5.5 | Paralysed |
| 0.5 | 100 | 1 | 5.8 | Paralysed |
| 5.0 | 1 | 2 | 5.5 | 2.6 |
| 5.0 | 1 | 3 | 5.5 | 5.8 |
| 5.0 | 1 | 4 | 6.0 | 0 |
| 1.0 | 10 | 1 | 6.0 | 6.0 |
| 1.0 | 10 | 2 | 5.6 | -0.3 |
| 1.0 | 15 | 2 | 5.5 | -0.8 |
| 1.0 | 20 | 2 | 5.8 | 1.5 some paralysis |
| 1.0 | 15 | 3 | 5.8 | -3.5 |

**Supplementary Table 1. 1 μL of 15% benzaldehyde (BZ) in ethanol for 3 hours induces learned avoidance.** Table shows the combinations of volume, concentration and duration tested for aversive conditioning. The optimal combination is shaded in grey

| **Strain name** | **Genotype** | **Reference** |
|---|---|---|
| N2 | N2 Bristol | CGC |
| CB4088 | *him-5(e1490)* | CGC |
| BAR109/<br>PT2248 | *pdf-1(tm1996)*III; *him-5(e1490)*V | This study<br>Barrios *et al.,* 2012 |
| PT2177 | *pdf-2(tm4393)* X; *him-5 (e1490)* V | Barrios *et al.,* 2012 |
| PT2159 | *pdf-2(tm4393)*X; *pdf-1(tm1996)*III;*him-5(e1490)*V | Barrios *et al.,* 2012 |
| BAR155 | *him-5 (e1490)*V; *pdf-1(tm1996)*III; *syb1393[AB01(pLAU-1-EG6699)MosCi LoxP::pdf-1cDNA::SL2::gfp::loxP::rfp]*II | This study |
| BAR167 | *him-5 (e1490)* V; *pdf-1(tm1996)*III; *syb1393* II*; oleEx39[grl-2::Cre+cc::rfp]* | This study |

| | | |
|---|---|---|
| BAR169 | *Him-5 (e1490)* V; *pdf-1(tm1996)*III;*syb1393* II;*kyEx4983 [sra-11::nCre(35ng/uL), myo-3::mCherry (5ng/uL)]* | This study |
| BAR170 | *him-5 (e1490)* V; *pdf-1(tm1996)*III;*syb1393* II;*kyEx4990[ceh-17::nCre(40ng/uL), myo-3::mCherry(5ng/uL)]* | This study |
| BAR187 | *pdf-2(tm4393)*X;*pdf-1(tm1996)*III;*him-5(e1490)*V;*syb1393* II | This study |
| BAR188 | *Pdf-2(tm4393)*X;*pdf-1(tm1996)*III;*him-5(e1490)*V;*syb1393* II*; oleEx39[grl-2(p)::Cre(30ng/µl)+cc::RFP(30ng/µl)]* | This study |
| BAR189 | *pdf-2(tm4393)*X;*pdf-1(tm1996)*III;*him-5(e1490)*V;*syb1393* II*; kyEx4983[sra-11::nCre (35ng/ul), myo-3::mCherry (5ng/ul)]* | This study |
| BAR108 | *pyIs500[(p)odr-3::gfp::egl-4+(p)odr-1::dsRed1+unc-122::GFP]*V;*him-5(e1490)*V | Lee *et al.,* 2010<br>This study |
| BAR224 | *him-5(e1490)*V; *oleEx80 [ttx-3::GCaMP6f::sl2::rfp (5ng/uL)]* | This study |
| BAR227 | *him-5(e1490)*V; *pdf-1(tm1996)*III; *oleEx80 [ttx-3::GCaMP6f::sl2::rfp (5ng/uL)]* | This study |
| BAR184 | *him-5(e1490)*V;*oleEx64[inx-1::GCaMP6f::sl2::rfp (15ng/ul)+ ttx-3::GCaMP6f::sl2::rfp (15ng/uL)]* | This study |
| BAR269 | *him-5(e1490)*V; *oleEx103[glr-3::GCaMP6f::sl2::rfp (30ng/uL)]* | This study |
| BAR253 | *him-5(e1490)*V;*oleEx95[tdc-1::GCaMP6f::sl2::rfp (10ng/uL)]* | This study |
| BAR122 | *pdfr-1(tm4457)*III;*him-5(e1490)*V;*myEx709[Ppdfr-1(3kb)::pdfr-1cDNAisofb+Ppdfr-1(3kb)::pdfr-1cDNAisofd+unc-122::GFP]* | This study |

| | | |
|---|---|---|
| BAR273 | *pdfr-1(tm4457)*III; *him-5(e1490)*V; *oleIs3[pdfr-1::LoxP::inv pdfr-1cDNAisofD::sl2::GFP::LoxP(10ng/µl)+ unc122::GFP(10ng/µl)]; oleIs5[rab-3::nCre (3ng/ul)+unc-122::RFP (30 ng/ul)]* | This study |
| BAR176 | *pdfr-1(tm4457)*III; *him-5(e1490)*V; *oleIs3* | This study |
| BAR206 | *pdfr1(tm4457)*III; *him-5(e1490)*V; *oleIs 3; oleEx72[tdc-1::nCre (20 ng/uL pSF178)+glr-3::nCre (50 ng/uL pSF179) + ttx-3::nCre (5 ng/uL pLAU 11)+ unc122::RFP(25ng/uL)]* | This study |
| PT2547 | *pdfr-1(tm4457)*III; *him-5(e1490)*V | Barrios *et al.,* 2012 |
| BAR196 | *lite-1(ce314)*X;*him-5(e1490)*V;*oleEx58[pdf-1::Campari (25 ng/µl) + cc::GFP (30ng/µl)]* | This study |
| BAR119 | *pdf-1(tm1996)*III;*him-5(e1490)*V; *oleEx39[grl-2(p)::Cre(30ng/µl)+cc::RFP(30ng/µl)]* | This study |
| CB369 | *unc-51(e369)* | CGC |
| BAR215 | *pdfr-1(tm4457)*III; *him-5(e1490)*V;*oleIs5* | This study |

**Supplementary Table 2.** Strains used in this study. CGC: Caenorhabditis Genetic Centre.

| Primer sequence (5'- 3') | Purpose | DNA Template |
|---|---|---|
| TTCCGGGGCGTTTTAAAAGTGG | *pdf-1(tm1996)* genotyping | *C.elegans* Genomic DNA |
| GTATGCGGACATCTTTTGTCTGC | *pdf-1(tm1996)* genotyping | *C.elegans* Genomic DNA |
| TGTGGTCAGAGTTCGTGTCA | *pdf-1(tm1996)* genotyping | *C.elegans* Genomic DNA |
| TAAAGGCAGTCAAGCTCGAAA | *pdf-1(tm1996)* genotyping | *C.elegans* Genomic DNA |
| CGGTTTTGTGGTAGCCTCAGTC | *pdf-2* genotyping | *C.elegans* Genomic DNA |
| CTAATTACGGATCCTTGTGGGCG | *pdf-2* genotyping | *C.elegans* Genomic DNA |

| | | |
|---|---|---|
| CGAGTGCTCTACAGTAAAGTTCGAG | *pdfr-1 (tm4457)* genotyping | *C.elegans* Genomic DNA |
| caaacaactgttgacacatcctgc | *pdfr-1 (tm4457)* genotyping | *C.elegans* Genomic DNA |
| **acgtcaccggttctagatacctagg**CGTGACGGGACACTCCTTATC | Lox-Ppdf-1::pdf-1-sl2-gfp Lox insert amplification | PSF-153 (Flavell *et al.,* 2013) |
| **ggctacgtaatacgactcacttaag**CTAGTAGGAAACAGTTATGTTTGGTATATTG | Lox-Ppdf-1::pdf-1-sl2-gfp Lox insert amplification | PSF-153(Flavell *et al.,* 2013) |
| CTTAAGTGAGTCGTATTACGTAGCC | *Mos ttTi5605* backbone amplification | PQL123 |
| CCTAGGTATCTAGAACCGGTG | *Mos ttTi5605* backbone amplification | PQL123 |
| **atgcgcggccgcactgactg**GCTTCAGTATTCATTTGTTCTTTTATAACTTCTC | *grl-2* promotor amplification (for *grl-2::nCre* assembly) | *C. elegans* Genomic DNA |
| **aatcccggggatcctctaga**ACTTGGTATAATTATGAAACTAACGAATATTGAATTTG | *grl-2* promotor amplification (for *grl-2::nCre* assembly) | *C. elegans* Genomic DNA |
| TCTAGAGGATCCCCGGGATTG | *nCre* backbone amplification | pEM-3 (Addgene) |
| CAGTCAGTGCGGCCGCGCAT | *nCre* backbone amplification | pEM-3 (Addgene) |
| **atgcgcggccgcactgactg**GCGAGTTTTGACTGGCTTTC | *rab-3* promotor amplification (for *rab-3::nCre* assembly) | rab-3::NLS::YFP (Ines Carrera lab) |
| **ctcttcttcttgggtgcgcccat**TTTTTCTGAGCTCGGTACC | *rab-3* promotor amplification (for *rab-3::nCre* assembly) | rab-3::NLS::YFP (Ines Carrera lab) |
| ATGGGCGCACCCAAGAAG | *nCre* backbone amplification | pEM-3 (Addgene) |
| CAGTCAGTGCGGCCGCGC | *nCre* backbone amplification | pEM-3 (Addgene) |
| **atgcgcggccgcactgactg**GTGAGCATCTATTACAGC | *ttx-3* promotor amplification (for *ttx-3::nCre* assembly) | pAB2 (ttx-3::GCaMP6f::SL2-RFP) |
| **ttcttcttgggtgcgcccat**TTTTTCTACCGGTACCCTC | *ttx-3* promotor amplification (for *ttx-3::nCre* assembly) | pAB2 (ttx-3::GCaMP6f::SL2-RFP) |

| | | |
|---|---|---|
| ATGGGCGCACCCAAGAAGAAGAG | *nCre* backbone amplification | pEM-3 (Addgene) |
| CAGTCAGTGCGGCCGCGC | *nCre* backbone amplification | pEM-3 (Addgene) |
| GCTAGCCAAGGGTCCTCC | Backbone and *inx-1* promotor amplification (for *inx-1::GCaMP6f::SL2-RFP* assembly) | pNP472 (Pokala *et al.*, 2014) |
| ATTCGTAGAATTCCAACTGAG | Backbone and *inx-1* promotor amplification (for *inx-1::GCaMP6f::SL2-RFP* assembly) | pNP472 (Pokala *et al.*, 2014) |
| **caggaggacccttggctagc**ATGGTCGACTCATCACGTCG | GCaMP6f-SL2-RFP amplification (for *inx-1::GCaMP6f::SL2-RFP* assembly) | pLR306 (Rene García Lab) |
| **tcagttggaattctacgaat**TTAGGCGCCGGTGGAGTG | GCaMP6f-SL2-RFP amplification (for *inx-1::GCaMP6f::SL2-RFP* assembly) | pLR306 (Rene García Lab) |
| ATTCGTAGAATTCCAACTGAG | Backbone and *ttx-3* promotor amplification (for *ttx-3::GCaMP6f::SL2-RFP* assembly) | ttx-3 in Ppd9575 (Hobert lab) |
| TTTTTCTACCGGTACCCTC | Backbone and *ttx-3* promotor amplification (for *ttx-3::GCaMP6f::SL2-RFP* assembly) | ttx-3 in Ppd9575 (Hobert lab) |
| **ggagggtaccggtagaaaaa**ATGGTCGACTCATCACGTCG | GCaMP6f-SL2-RFP amplification (for ttx-3*::GCaMP6f::SL2-RFP* assembly) | pLR306 (Rene García Lab) |
| **tcagttggaattctacgaat**TTAGGCGCCGGTGGAGTG | GCaMP6f-SL2-RFP amplification (for ttx-3*::GCaMP6f::SL2-RFP* assembly) | pLR306 (Rene García Lab) |
| ATGGTCGACTCATCACGTCGTAAGTGG | GCaMP6f-SL2-RFP backbone amplification (for *tdc-1::GCaMP6f::SL2-RFP* assembly) | pAB1 (*inx-1::GCaMP6f::SL2-RFP*) |
| GGCCGGCCCAGTCAGTGC | GCaMP6f-SL2-RFP backbone | pAB1 |

| | | |
|---|---|---|
| | amplification (for *tdc-1::GCaMP6f::SL2-RFP* assembly) | (*inx-1::GCaMP6f::SL2-RFP*) |
| **ccgcactgactgggccggcc**CACCTAACTTCGTCGGTCAAC | *tdc-1* promotor amplification (for *tdc-1::GCaMP6f::SL2-RFP* assembly) | PJM11 (Barrios Lab) |
| **cgacgtgatgagtcgaccat**GGTACCCTCCAAGGGTCC | *tdc-1* promotor amplification (for *tdc-1::GCaMP6f::SL2-RFP* assembly) | PJM11 (Barrios Lab) |
| GGCGCGCCTCTAGAGGATCC | GCaMP6f-SL2-RFP backbone amplification (for *glr-3::GCaMP6f::SL2-RFP* assembly) | pAB1 |
| GGCCGGCCCAGTCAGTGC | GCaMP6f-SL2-RFP backbone amplification (for *glr-3::GCaMP6f::SL2-RFP* assembly) | pAB1 |
| **ccgcactgactgggccggcc**TCGGAAATGCGGAAGTTC | *glr-3* promotor amplification (for *glr-3::GCaMP6f::SL2-RFP* assembly) | pSF179 (Flavell *et al.,* 2013) |
| **ggatcctctagaggcgcgcc**ATGTTAATAGCAAATATTGAAGATTCTAAAC | *glr-3* promotor amplification (for *glr-3::GCaMP6f::SL2-RFP* assembly) | pSF179 (Flavell *et al.,* 2013) |

**Supplementary Table 3:** Primers used in this study. Primers used to build the same construct are entered consecutively and highlighted as a group in white or grey. In primers used for Gibson assembly, the overlapping region between insert and backbone vector is highlighted in bold.

| Plasmid name | DNA construct | Reference |
|---|---|---|
| pLAU-1 | *Mos ttTi5605-pdf-1::lox pdf-1-sl2-GFPlox RFP unc-119 MosttTi5605* | This study |
| pLAU-2 | *grl-2::nCre* | This study |
| pLAU-4 | *rab-3::nCre* | This study |
| pLAU-9 | *glr-3::GCAMP6f-sl2-RFP* | This study |
| pLAU-10 | *tdc-1::GCAMP6f-sl2-RFP* | This study |
| pLAU-11 | *ttx-3::nCre* | This study |
| pSF134 | *Pdfr-1 distal floxed pdfr-1 iso d* | Flavell *et al.,* 2013 |

| | | |
|---|---|---|
| pSF153 | *pdf-1::lox pdf-1-sl2-GFPlox RFP* | Flavel et al 2013 |
| pSF178 | *tdc-1::nCre* | Flavel et al 2013 |
| pSF179 | *glr-3::nCre* | Flavel et al 2013 |
| pEM3 | *ncs-1::nCre* | Addgene |
| pRTAB1 | *pdf-1::CAMPARI* | This study |
| pAB1 | *inx-1::GCaMP6f-sl2-RFP* | This study |
| pAB2 | *ttx-3::GCaMP6f-sl2-RFP* | This study |
| pQL123 | *Mos ttTi5605 -unc-119-MosttTi5605* | Queelim Ch'ng Lab |

**Supplementary Table 4:** Plasmids used in this study
